# Comprehensive molecular characterization of pediatric treatment-induced high-grade glioma: A distinct entity despite disparate etiologies with defining molecular characteristics and potential therapeutic targets

**DOI:** 10.1101/809772

**Authors:** John DeSisto, John T. Lucas, Ke Xu, Andrew Donson, Tong Lin, Bridget Sanford, Gang Wu, Quynh T. Tran, Dale Hedges, Chih-Yang Hsu, Gregory T. Armstrong, Michael Arnold, Smita Bhatia, Patrick Flannery, Rakeb Lemma, Lakotah Hardie, Ulrich Schüller, Lindsey M. Hoffman, Kathleen Dorris, Jean Mulcahy Levy, Todd C. Hankinson, Michael Handler, Arthur Liu, Nicholas Foreman, Rajeev Vibhakar, Kenneth Jones, Sariah Allen, Jinghui Zhang, Suzanne J. Baker, Thomas E. Merchant, Brent A. Orr, Adam L. Green

**Affiliations:** Morgan Adams Foundation Pediatric Brain Tumor Research Program, University of Colorado School of Medicine, Aurora, CO; Department of Radiation Oncology, St. Jude Children’s Research Hospital, Memphis, TN; Computational Biology, St. Jude Children’s Research Hospital, Memphis, TN; Biostatistics, St. Jude Children’s Research Hospital, Memphis, TN; Pathology, St. Jude Children’s Research Hospital, Memphis, TN; Epidemiology and Cancer Control, St. Jude Children’s Research Hospital, Memphis, TN; Developmental Neurobiology, St. Jude Children’s Research Hospital, Memphis, TN; Childhood Cancer Survivor Study; Department of Pathology, Nationwide Children’s Hospital, Columbus, OH; Division of Hematology and Oncology, University of Alabama at Birmingham; Children’s Hospital Colorado, Aurora, CO; Institute of Neuropathology and Department of Pediatric Hematology and Oncology, University Medical Center; Children’s Cancer Center, Hamburg, Germany

## Abstract

Treatment-induced high-grade gliomas (TIHGGs) are an incurable late complication of cranial radiation therapy or combined radiation/chemotherapy used to treat pediatric cancer. We assembled a cohort of 33 TIHGGs from multiple institutions. The primary antecedent malignancies were medulloblastoma, acute lymphoblastic leukemia, astrocytoma, and ependymoma. We performed methylation profiling, RNA-seq, and genomic sequencing (whole-genome or whole-exome) on TIHGG samples. Methylation profiling revealed that TIHGGs cluster primarily with the pediatric receptor tyrosine kinase I subtype (26/31 samples). Common TIHGG copy-number alterations include Chromosome (Ch.) 1p loss/1q gain, Ch. 4 loss, Ch. 6q loss, and Ch. 13 and Ch. 14 loss; focal alterations include *PDGFRA* and *CDK4* gain and loss of *CDKN2A* and *BCOR*. Relative to *de novo* pediatric high-grade glioma (pHGG), *BCOR* loss (p=0.004) and *CDKN2A* loss (p=0.005) were significantly increased. Transcriptomic analysis identified two distinct TIHGG subgroups, one with a lesser mutation burden (0.12 mut/Mb), Ch. 1p loss/1q gain (5/6 samples), and stem cell characteristics, and one with a greater mutation burden (1.08 mut/Mb, p<0.0002), depletion of DNA repair pathways, and inflammatory characteristics. We observed increased chromothripsis in TIHGG versus pHGG (67% vs. 31%, p=0.036), which was associated with extrachromosomal circular DNA-mediated amplification of *PDGFRA* and *CDK4*. *In vitro* drug screening in one primary, patient-derived TIHGG cell line from each expression subgroup identified microtubule inhibitors/stabilizers, DNA-damaging agents, MEK inhibition, and, in the inflammatory subgroup, proteasome inhibitors as potentially effective therapies. This study provides a comprehensive molecular profile of TIHGG, including mechanistic insights to TIHGG oncogenesis, and identifies potentially effective therapeutic modalities for further investigation.

## Introduction

Treatment-induced high-grade gliomas (TIHGGs) are pediatric high-grade astrocytomas with an estimated incidence of 3% following cranial radiotherapy^1^. The highest cumulative incidence of TIHGG is in survivors of medulloblastoma, ependymoma, and leukemia^1^. TIHGG represents a rare but significant cause of late mortality in survivors of childhood cancer^2^, as few therapeutic options are available at diagnosis^3, 4^. Genomic analysis of TIHGG has been restricted to only a few small cohorts, but early evidence suggests that, in comparison with *de novo* pediatric high-grade gliomas (pHGG), TIHGG may be enriched for recurrent copy-number alterations and for amplification of *PDGFRA* and other potential activators of the MAP kinase pathway^5, 6^. *TP53* mutations, gain of 1q, deletion of *CDKN2A*, and overexpression of *ERBB3* and *SOX10* may also be more frequent^5, 7^. Similarly, alterations in treatment-related second cancers outside the brain show recurrent genomic copy-number variations, some of which contain genes that modify tumor latency *(NF2*, *CHEK2*, and *RET* amplification)^8,9,10^. These divergent oncogenic programs are spread over multiple histologies, limiting inference regarding potential therapeutic options.

In this study, we present an evaluation of the largest TIHGG cohort assembled to date, using comprehensive clinical and molecular analyses. We conducted a clinical review of each case, including the initial tumor location and type, treatment received, and the resulting TIHGG location and histology, followed by DNA methylation analysis using 450/850k arrays, genomic sequencing of TIHGG and matched blood samples, and RNA-seq. We assemble a comprehensive molecular profile of TIHGG, enabling us to identify several previously unknown characteristics of the disease, and propose a method of grouping TIHGG tumors phenotypically based upon a combination of genomic alterations and gene expression characteristics. Our findings differentiate TIHGG from pHGG and provide rationale for alternative therapeutic approaches, which we explore in *in silico* and *in vitro* pre-clinical therapeutic screening studies.

## Results

### Patient characteristics

The median age at diagnosis of first cancer within the TIHGG cohort was 6.5 years (range, 0.16-19). The initial diagnoses included medulloblastoma (N=12; 36%), acute lymphoblastic leukemia (ALL) (N=10; 30.3%), astrocytoma (N=3; 9%), ependymoma (N=2; 6%), germinoma (N=2; 6%), lymphoma (N=1; 3%), craniopharyngioma (N=1; 3%), neuroblastoma (N=1; 3%), and ganglioglioma (N=1; 3%) (Sup. Table 1). Two patients had known mismatch repair deficiencies resulting from a heterozygous germline loss of *MSH2* in one case and *PMS2* in the other (Case 2, Case 32). Both were included in the study because each had therapeutic radiation exposure to the cranium with sufficient latency (11 and 8 years) to meet the TIHGG diagnostic criteria. All cases had ionizing radiation exposure, except for one case with a prior history of high-dose chemotherapy exposure for treatment of high-risk neuroblastoma (further details on clinical histories are discussed in Methods). The individual clinical histories for the TIHGG cohort with extended clinical information are presented in Sup. Fig. 1.

Complete clinical-pathologic data regarding treatment history were available in 76% of cases (Fig. 1A (showing analyses/information available for the entire cohort), Sup. Fig. 1, Sup. Table 1, 2). Patients received total body irradiation (TBI) (N=1, 3%), cranial irradiation (N=6, 18%), craniospinal irradiation (N=9, 24%), or focal brain irradiation (N=10, 29%) to treat the initial malignancy (Fig. 1B). One patient received radiotherapy to the abdomen (Case 44). Radiotherapy field extent was unknown in 24% (N=7) of cases. The indications for radiotherapy were adjuvant (N=12, 36%), definitive (N=1, 3%), and salvage (N=9, 26.5%). Radiotherapy technique was 2D (N=13, 38.2%) or 3D conformal radiation therapy (N=8, 23.5%). Technique was not known in 38% (N=13) of cases. The location of the TIHGG relative to the initial radiotherapy field was in-field but outside the high-dose region in 24% (N=8) of evaluable cases and in the high-dose (prescribed dose) region in 52% of cases (N=17).

**Figure 1.**
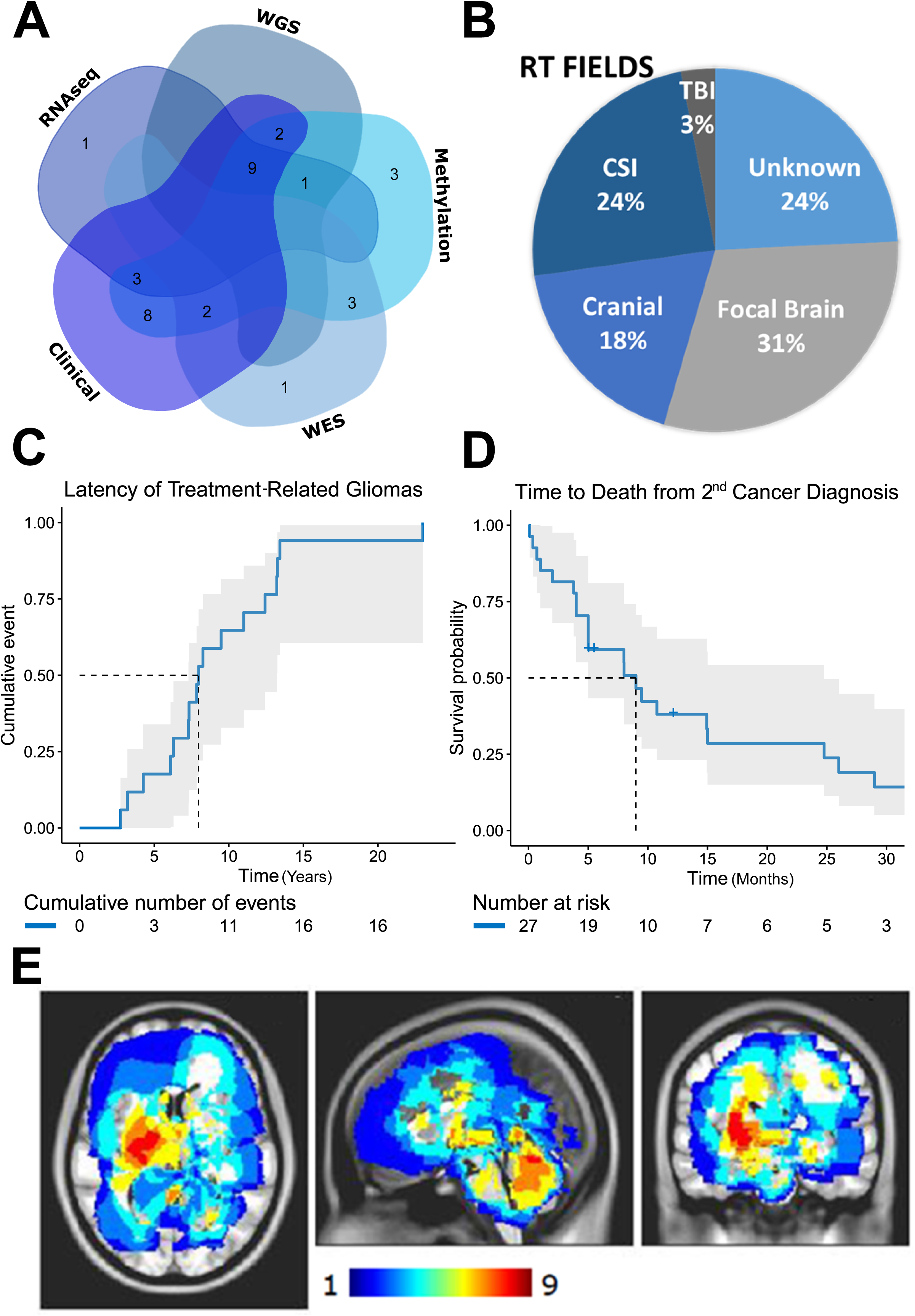
TIHGG cohort characteristics. A) Venn diagram illustrating the number of cases with each type of historical data/molecular analysis. B) Radiotherapy field type used for initial cancer. C) Time to diagnosis of TIHGG from initial cancer diagnosis. D) Time to death following TIHGG diagnosis. *Two patients survived beyond 35 months but were deceased at study closure. E) Anatomic location of TIHGG. Color scale indicates the number of cases anatomically overlapping at each point in space.

The median time between initial treatment and TIHGG diagnosis was 8.0 years (95% CI 7.3-13.2 years) (Fig. 1C). The median age at the time of diagnosis of TIHGG was 13.6 years (9-26 years) (Sup. Table 1). TIHGG tumor grades were WHO grade III (N=3, 9%), grade IV (N=12, 55%), and pHGG not otherwise specified (N=12, 36%) (Sup. Table 1). Assessment by methylation array identified *MGMT* promoter methylation in 8 of 31 (25.8%) cases in which methylation status was evaluated during the clinical course of treatment.

Salvage therapy characteristics for TIHGG were available in 38% of cases (N=13) and included resection (N=3, 23.1%), repeat radiotherapy (N=12, 92.3%), and/or chemotherapy (N=4, 30.7%) (Sup. Table 2). The median radiotherapy dose for the second course was 54 Gy (range 30-59.4 Gy). At the time of this analysis, all TIHGG patients with known outcome were deceased (N=26), and 7 had unknown clinical status (Fig. 1D). The median survival in the TIHGG cohort was 9 months (95% CI 5-24.7 months) (Fig. 1D, Sup. Table 2).

The use of radiotherapy, chemotherapy, and/or resection did not substantially improve treatment outcomes, although a paucity of details available regarding therapy limited power for this analysis (Sup. Table 2). These outcomes are consistent with prior published clinical series of TIHGG^4, 8^.

### Voxel-wise lesion mapping of TIHGG relative to *de novo* pHGG

An MRI was available at diagnosis in 21 TIHGG cases (Sup. Fig. 2A). In cases for which location information was available, TIHGGs were located in the infratentorial brain (posterior fossa) (N=11, 32.3%), supratentorial brain (N=12, 35.3%), supra- and infratentorial brain (N=1, 3%), and spine (N=1, 3%) (Fig. 1E, Sup. Fig. 2B). An additional cohort of 84 *de novo* pHGG was used for comparison. Compared to a cohort of 84 de novo pHGG, TIHGG tumors were more likely to involve the frontal and occipital lobe, and posterior fossa (cerebellum and inferior brainstem) regions relative to primary pHGG (Fig. 1E, Sup. Fig. 2B).

### TIHGGs cluster into a specific methylation group

We began our molecular analysis using DNA methylation profiling because methylation data (or a DNA sample sufficient to conduct methylation profiling) were available for the greatest number of TIHGG samples in our cohort. We first analyzed the methylation data using unsupervised clustering of 34 TIHGG samples from 31 tumors against a combined reference cohort containing pediatric and adult CNS malignancies^11, 12^. Twenty-five of 31 cases clustered among a group of epigenetically similar tumors consisting of *H3K27M*-negative midline gliomas and gliomas in the pediatric receptor tyrosine kinase I (pedRTK I) subgroup (Fig. 2A-C and Sup. Fig. 3A-C)^13^. TIHGGs were not readily distinguishable by methylation from IDH wt, GBM-MID, and pedRTK I cases from the Capper and Korshunov reference cohorts^11^, so the combined group of TIHGG cases is designated as “pedRTK I” going forward, reflecting their closer overall proximity to the centroids of the pedRTK I reference cases (Sup. Table 3). The remaining six TIHGGs clustered with the GBM K27M (N=1), control cerebellum (N=2), anaplastic pilocytic astrocytoma (N=1), HGNET-MN1 (N=1), and pleomorphic xanthoastrocytoma (N=1) methylation groups. We confirmed the initial classification results using hierarchical and supervised methods (Sup. Table 4 and Sup. Table 5). No trend was noted associating initial cancer diagnosis with subsequent TIHGG methylation class (Sup. Fig. 4A)^14^.The latency and survival of TIHGG tumors classified as pedRTK I did not differ significantly from that of those classified with other subclasses (Sup. Fig. 4B-C). The odds of being assigned to the pedRTK I subgroup in the TIHGG cohort were 29.2 times the odds of being assigned to that subgroup in the *de novo* pHGG cohort (95% CI 9.15-92.9, p<0.0001) (Sup. Fig. 5A, Sup. Table 6). Survival of the TIHGG-pedRTK I was inferior to pHGG pedRTK I (median survival 9 months, [95% CI, 5-26] vs. 16 months [95% CI, 10-not reached, p=0.00021] (Sup. Fig. 5B).

**Figure 2.**
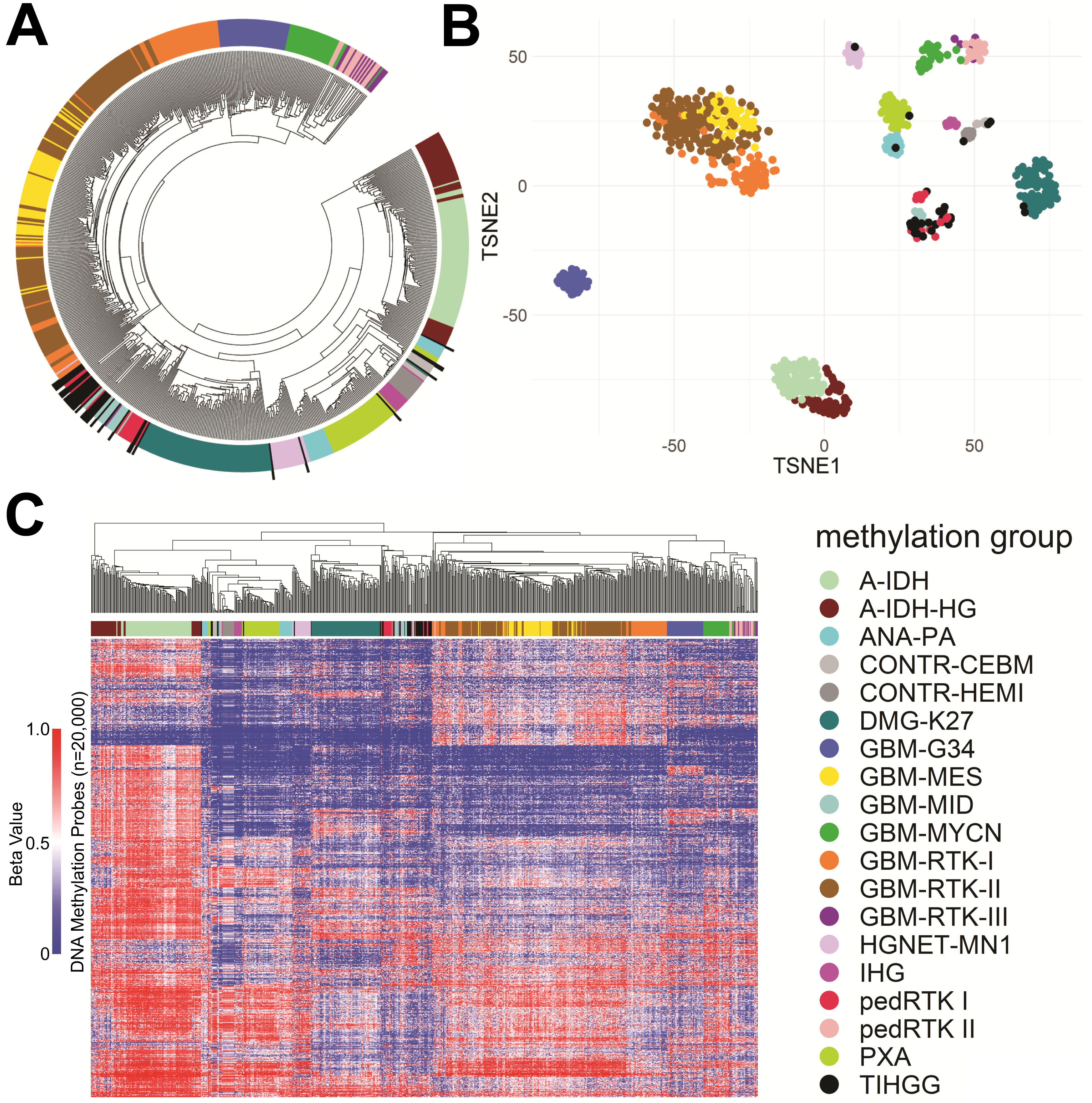
Methylation-based classification of TIHGG. A) Circular dendrogram indicating location of TIHGG (black bars) relative to diagnoses. B) Localization of TIHGG relative to other CNS cancers in t-distributed stochastic neighboring (t-SNE) space. C) Dendrogram and heatmap of TIHGG cases clustered against other pediatric brain tumor entities. A-IDH= Astrocytoma, subclass IDH-mutant; A-IDH-HG= High-Grade Astrocytoma, subclass IDH-mutant; ANA PA = Anaplastic Pilocytic Astrocytoma, CONTR-CEBM= Control Cerebellum; CONTR-HEMI= Control Cerebral Cortex; DMG-K27 = Diffuse Midline Glioma, H3 K27M-mutant; GBM-G34= Glioblastoma, subclass H3.3 p.G34R-mutant; GBM-MES= Glioblastoma, subclass mesenchymal; GBM-MID = Glioblastoma, IDH-Wild type, subclass midline; GBM-MYCN= Glioblastoma, subclass MYCN-amplified; GBM-RTK-I = Adult Glioblastoma, subclass RTK I; GBM-RTK-II= Adult Glioblastoma, subclass RTK II; GBM-RTK-II = Adult Glioblastoma, subclass RTK III; HGNET-MN1 = High Grade Neuroepithelial Tumor with MN1 alteration; IHG= Infantile High Grade Glioma; pedRTK I= Pediatric Glioblastoma, subclass RTK I; pedRTK II = Pediatric Glioblastoma, subclass RTK II; PXA = Pleomorphic Xanthoastrocytoma; TIHGG = Treatment-Induced High-Grade Glioma.

Alignment of TIHGGs with pedRTK I suggested the possibility that TIHGGs, despite their disparate clinical precursors, might share common molecular alterations, which we investigated by comparing copy-number alteration and transcriptomic profiles discussed in the following sections.

### TIHGGs show recurrent copy-number abnormalities

We next reviewed the methylation profiling data for large segment and focal copy-number alterations. Total copy-number variation was significantly increased in TIHGG when comparing *de novo* pHGG and TIHGG in the pedRTK I subgroup (Sup. Fig. 5C). Recurrent large segment copy-number alterations identified from methylation profiling included Ch.1p loss (3/31, 9.7%), Ch.1q gain (3/31, 9.7%), Ch.6q loss (5/31, 16.1%), Ch.13 loss (8/31, 25.8%), and Ch.14 loss (3/31, 9.7%) (Fig. 3A, Sup. Table 5, Sup. Fig. 6). Copy-number alterations in *PDGFRA* gain/amplification (15/31, 48.4%), *CDK4* amplification (5/31, 12.9%), *CDKN2A* loss (12/31, 38.7%), and *BCOR* loss (7/31, 22.6%) were also noted. These were increased relative to *de novo* pHGG for *PDGFRA* (p=0.0055) and *BCOR* (p=0.0043) (Sup. Table 7, Sup. Fig. 7). Changes in copy number of *PDGFRA, CDKN2A,* and *BCOR* were associated with an accompanying change in gene expression in cases with available RNA-Seq data (N=12, 34.3%) (Fig. 3B). When comparing total genomic copy-number variations by case, TIHGG favored loss over gain (Fig. 3C).

**Figure 3.**
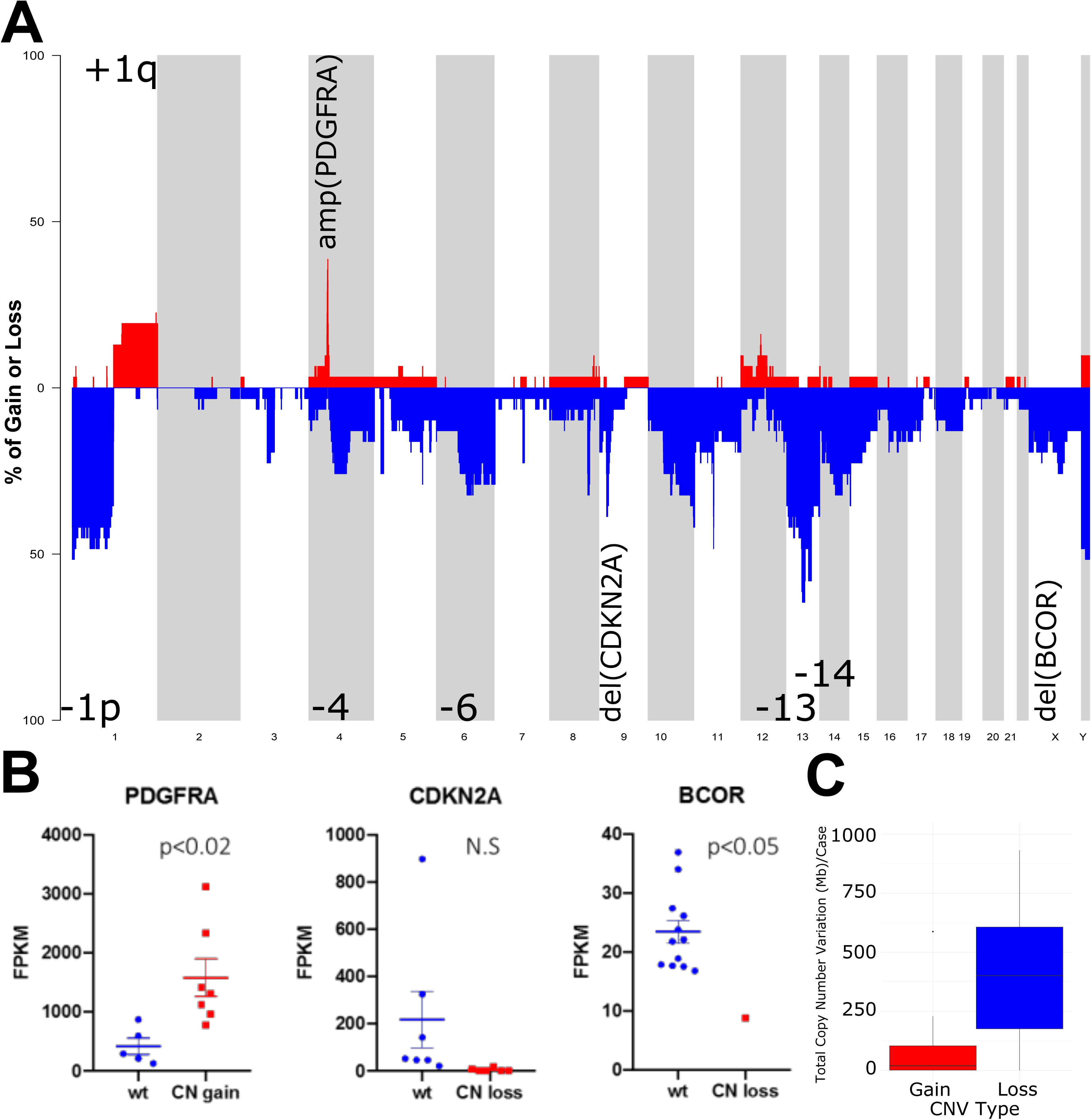
Frequent copy number alterations in TIHGG. A) Copy-number frequency plot. B) Expression-based confirmation of high frequency copy number alterations in *PDGFRα, CDKN2A,* and *BCOR;* mean and standard error of the mean (SEM) shown. C) Frequency of copy number gain and loss across all TIHGG; boxes show median and 1^st^/3^rd^ quartiles, with whiskers representing range limited to 1.5x the interquartile range from the box edge.

Overall, TIHGGs showed large segment genomic alterations in Ch.1 (1p loss/1q gain) and Ch.13 (loss), and focal alterations in *PDGFRA* (amplification)*, CDKN2A* (loss) and *BCOR* (loss). As further discussed below, where alterations were identified in both WES/WGS and methylation data, we obtained consistent results between the two methods. The *PDGFRA* amplification that we observed is consistent with the alignment of most TIHGGs with the pedRTK I methylation subclass^11^.

### TIHGG has a high frequency of chromothripsis and episomal amplification of oncogenes

To further investigate the potential sources of copy-number amplification, we evaluated the rate of chromothripsis in the TIHGG samples relative to that in primary non-brainstem and brainstem pHGG from Wu *et al.*^15^. We observed an increase in the frequency of chromothripsis in the TIHGG cohort (8/12 cases) relative to that in diffuse intrinsic pontine glioma (DIPG) (66.7% vs. 30.0%, p=0.048), non-brainstem pHGG (66.7% vs. 33.3%, p=0.091), and all primary pHGG combined (66.7% vs. 31.4%, p=0.036) (Fig. 4A, Sup. Table 8). Both intra-chromosomal (Case 29-dm2) rearrangements leading to extrachromosomal circular DNA (eccDNA) structures, as well as inter-chromosomal rearrangements (Case 29-dm1) leading to eccDNA structures, were observed in case 29 (Fig. 4B, D). *PDGFRA* and *CDK4* appeared to be amplified separately on the two eccDNA structures (Fig. 4B, D). Amplification was confirmed in selected cases by fluorescence *in situ* hybridization, which demonstrated a double minute pattern, a finding typical of episomal amplification (Fig. 4C)^16^. There was a trend toward an increased frequency of chromothripsis in cases with a coincident pathogenic *TP53* mutation (5/8 cases, 62.5%) relative to those with wild-type *TP53* (0/4 cases, 0%, p=0.08).

**Figure 4.**
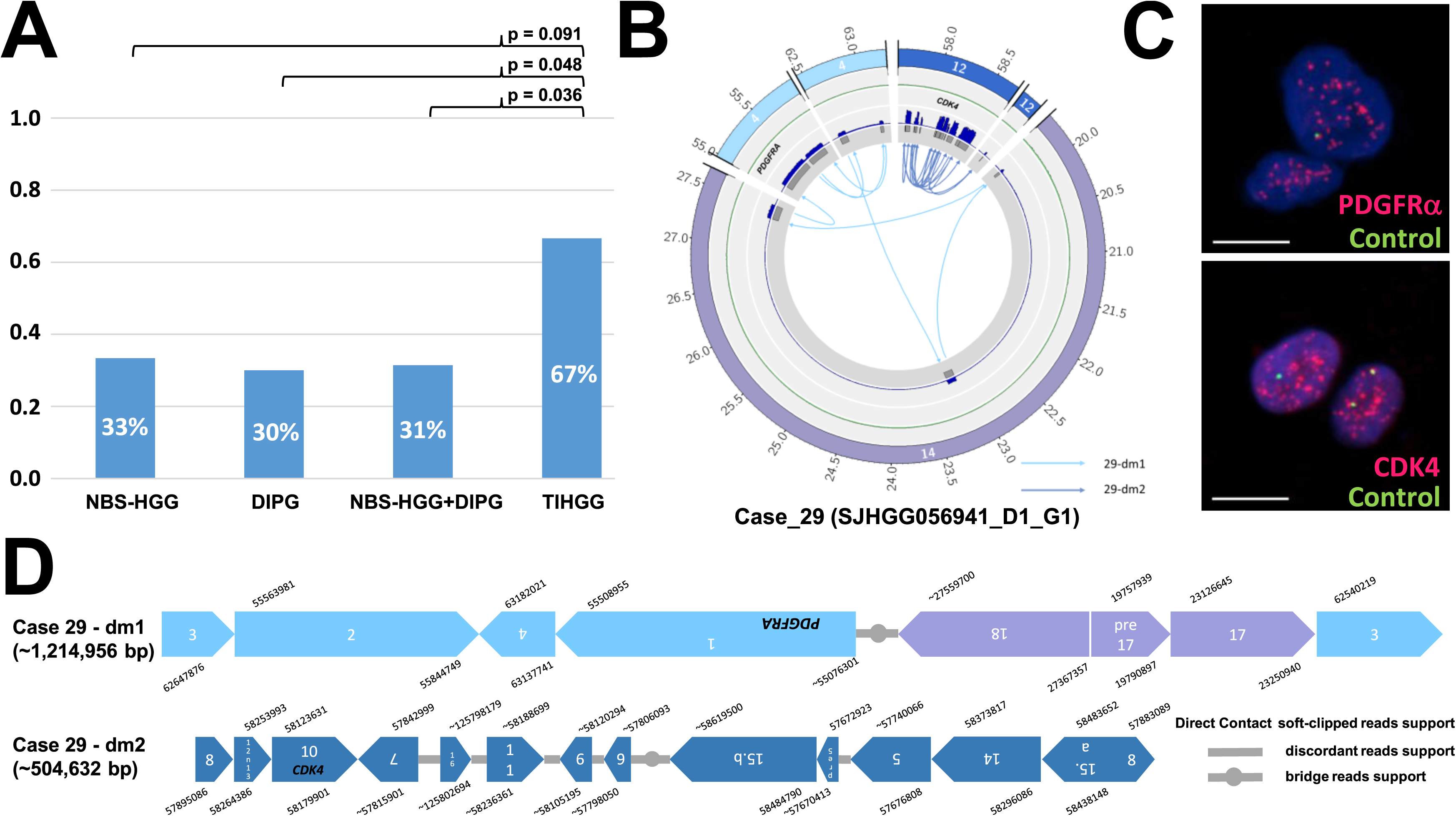
Frequency of chromothripsis and resultant eccDNA structures, oncogene incorporation and FISH analysis in Case_29. A) Frequency of chromothripsis in primary non-brainstem high-grade glioma (NBS-HGG), primary DIPG, TIHGG, and combined NBS-HGG and DIPG. B) Circos plot illustrating the intra- and inter-chromosomal rearrangements of chromosomes 4 and 14, resulting in dm1, and intra-chromosomal rearrangements in chromosome 12 resulting in dm2. Byproduct of chromothripsis events between chromosomes 4 and 12 (dm1) and intra-chromosomal rearrangements in chromosome 14 (dm2). The innermost circle shows the highly amplified CNA segments, as evidenced by their extremely high coverage in the tumor sample (blue circle) compared to that in the paired germline sample (green circle). The outermost circle shows the involved chromosomes. C) Two-color FISH experiment from Case_29 shows *PDGFRα* (upper panel) and *CDK4* amplification (lower panel) with diffuse punctate staining relative to the control centromeric DNA in each cell, suggesting the presence of multiple double-minutes. D) The predicted structures of the eccDNA dm1 and dm2 showing the directions of each of the joined CNA segments and their break point positions. “Bridge reads support” means SVs identified through common discordant reads support between two segment boundaries as described in Xu *et al*^16^.

In summary, we observed an elevated frequency of chromothripsis in TIHGG relative to *de novo* pHGG, which facilitated extrachromosomal amplification of known oncogenes (*PDGFRA, CDK4)*.

### TIHGGs show recurrent abnormalities in genes that regulate cell proliferation, growth, and cell cycle

In addition to copy-number variation, we also identified focal alterations in TIHGG using WES/WGS data. Eighteen TIHGG cases underwent whole-exome (WES) or whole-genome sequencing (WGS) (Fig. 5). Tier 1 mutations in the 5 non-hypermutator WES cases without matched germline data are shown in Sup. Table 9; Tier 1 mutations for the 9 non-hypermutator WGS cases with matched germline data are shown in Sup. Table 10. The most frequent focal alterations included *PDGFRA*, *CDKN2A*, *BCOR*, *NF1*, *TP53*, and *CDK4* (Fig. 5, Sup. Tables 7, 9, 10). Missense mutations were noted in *TP53* and *PDGFRA* in 7 of 8 and 4 of 15 cases with an observed alteration in *TP53* and *PDGFRA,* respectively. Nonsense and frameshift mutations of *NF1* occurred in 8 of 9 observed cases with *NF1* alterations. *MET* fusion gene products were identified from RNA-seq samples in three cases (Sup. Table 11). Structural variations and copy-number alterations from 10 WGS cases are shown in Sup. Tables 12 and 13.

**Figure 5.**
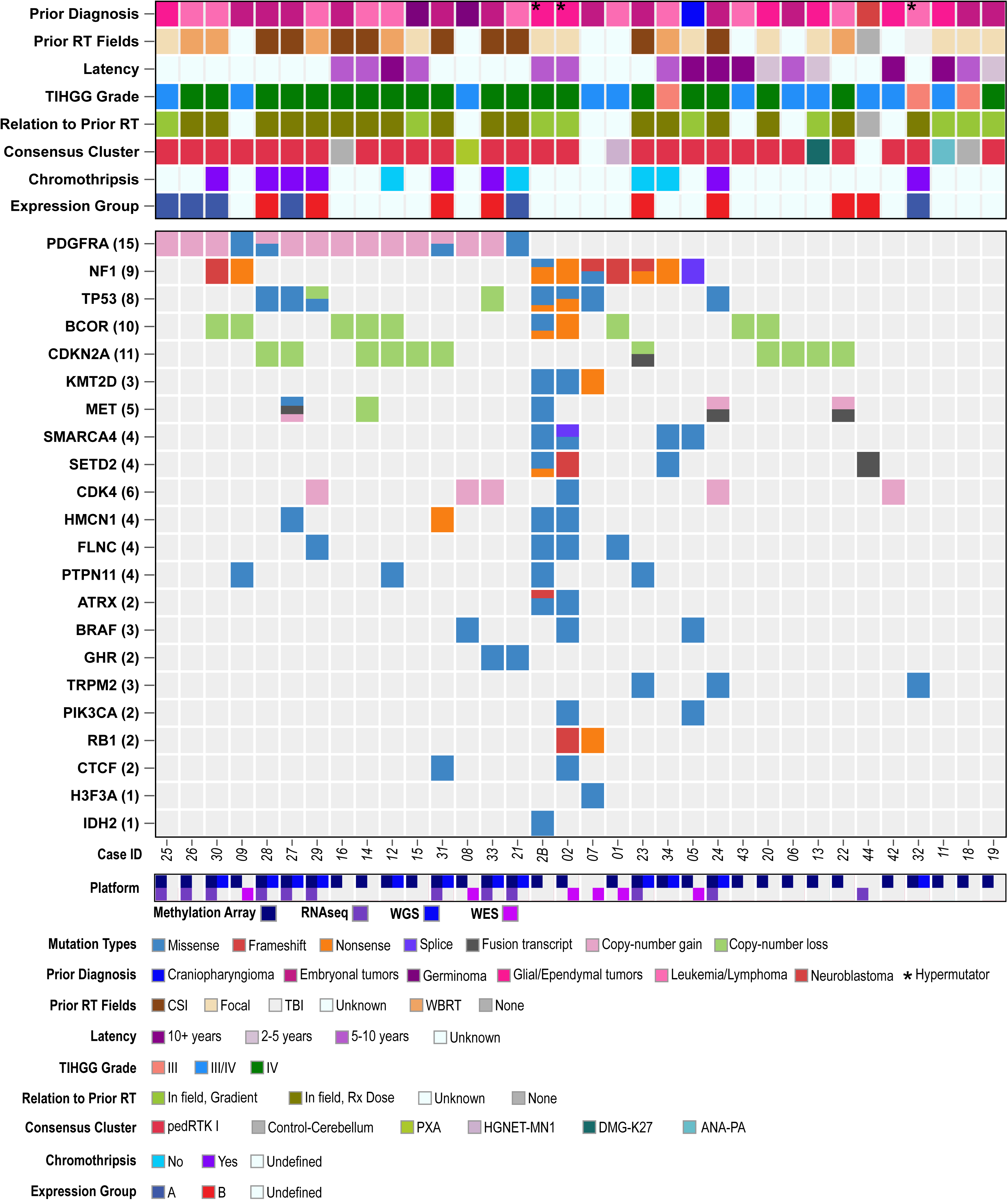
Recurrent molecular alterations in TIHGG. Oncoprint describing the clinical characteristics, histopathological features, methylation profile, tier1 mutations, genes affected by copy-number gain/loss, and fusion genes. ANA PA = Anaplastic Pilocytic Astrocytoma; CONTR-CEBM = Control Cerebellum; DMG-K27 = Diffuse Midline Glioma H3.3 K27M; GBM-MID = Glioblastoma IDH-wild type, subclass midline; HGNET-MN1 = High Grade Neuroepithelial Tumor with MN1 alteration; PXA = Pleomorphic Xanthoastrocytoma. Sequencing/array platforms for each case are also shown on the bottom row. The second sample for Case 2 is shown as Case 2B while the second sample for Case 14 and Case 12 are not shown but are annotated in Sup. Table 5.

### Compared to pHGG, TIHGG are enriched for specific somatic and germline abnormalities

We compared select recurrent alterations in the TIHGG cohort to *de novo* pHGG from the HERBY cohort across each cohort as a whole and among pedRTK I tumors (Sup. Fig. 8, Sup. Table 14)^14^. Statistically significant increases in *BCOR* loss (p=0.0004) and *CDKN2A* loss (p=0.05) were observed in TIHGG relative to pHGG (Sup. Table 7). No statistically significant differences were observed between the proportions of molecular alterations in pedRTK I TIHGG relative to pHGG – pedRTK I tumors, although power was limited by the sparse number of pedRTK I cases in the HERBY cohort (N=8). Of note, the frequency of *TP53* mutations in pedRTK I TIHGG cases was 43.8% (7/16) and 62% (5/8) in the *de novo* pHGG pedRTK I cases (p=0.68, Sup. Table 7). We observed a single *H3F3A* mutation (Fig. 5, Case 7) (1/19, 5.6%) in the TIHGG cohort, compared to 37.8% (28/74) with mutant *H3F3A* (p=0.0053) in the HERBY cohort (composed of non-brainstem pHGG).

We also compared variant frequency between the TIHGG cohort and pHGG cases in the HERBY cohort. Germline mutations in the TIHGG cohort were analyzed in 10 cases for which WGS data were available (Sup. Table 15). The total number of germline and somatic variants per megabase (Mb) were calculated and stratified by pHGG versus TIHGG and are compared in Sup. Fig. 9A. The number of variants per Mb was significantly higher in somatic coding regions of TIHGG DNA than in those of pHGG (median 0.000784 ± 0.000286 (range) variants/Mb vs. median 0.0007 ± 0.000245 (range) variants/Mb, p=0.031). The relative frequencies of base transitions were calculated per sample in pHGG and TIHGG (Sup. Fig. 9B). Germline and somatic coding base changes were not significantly different across base change type. Somatic noncoding region variants showed decreased A to C (median 14.4 ± 4.91 vs. 8.8 ± 4.1, p=0.007) and A to G (34.3 ± 9.67 vs. 20.2 ± 11.3, p=0.03), and increased C to A (6.79 ± 2.2 vs. 16.7 ± 5.47, p=0.004) base changes in TIHGG cases relative to *de novo* pHGG (Sup. Fig. 9B-C).

In summary, TIHGG have comparable burden of somatic and germline alterations to *de novo* pHGG. Frequent pathogenic alterations were noted in *PDGFRA, BCOR, CDK4, TP53,* and *NF1*.

### Transcriptomic analysis demonstrates that TIHGGs cluster into two subgroups that are separate from pHGG

Transcriptomic data (array and RNA-seq) were available on 14 TIHGG cases and 42 *de novo* glioblastoma (GBM) cases. TIHGG tumors clustered separately from *de novo* pediatric, infant, and adult GBM based on transcriptomic data and formed two distinct subgroups (A and B) containing 6 and 8 tumor samples respectively (Fig. 6A). A comparison of the transcriptomic and methylation-based clustering results showed that the two gene expression-based TIHGG subgroups were, with few exceptions, recapitulated in the methylation analysis (Fig. 6B). Our data thus suggest two distinct patterns of gene expression within the pedRTK I methylation-based subgroup into which most of the TIHGGs cluster.

**Figure 6.**
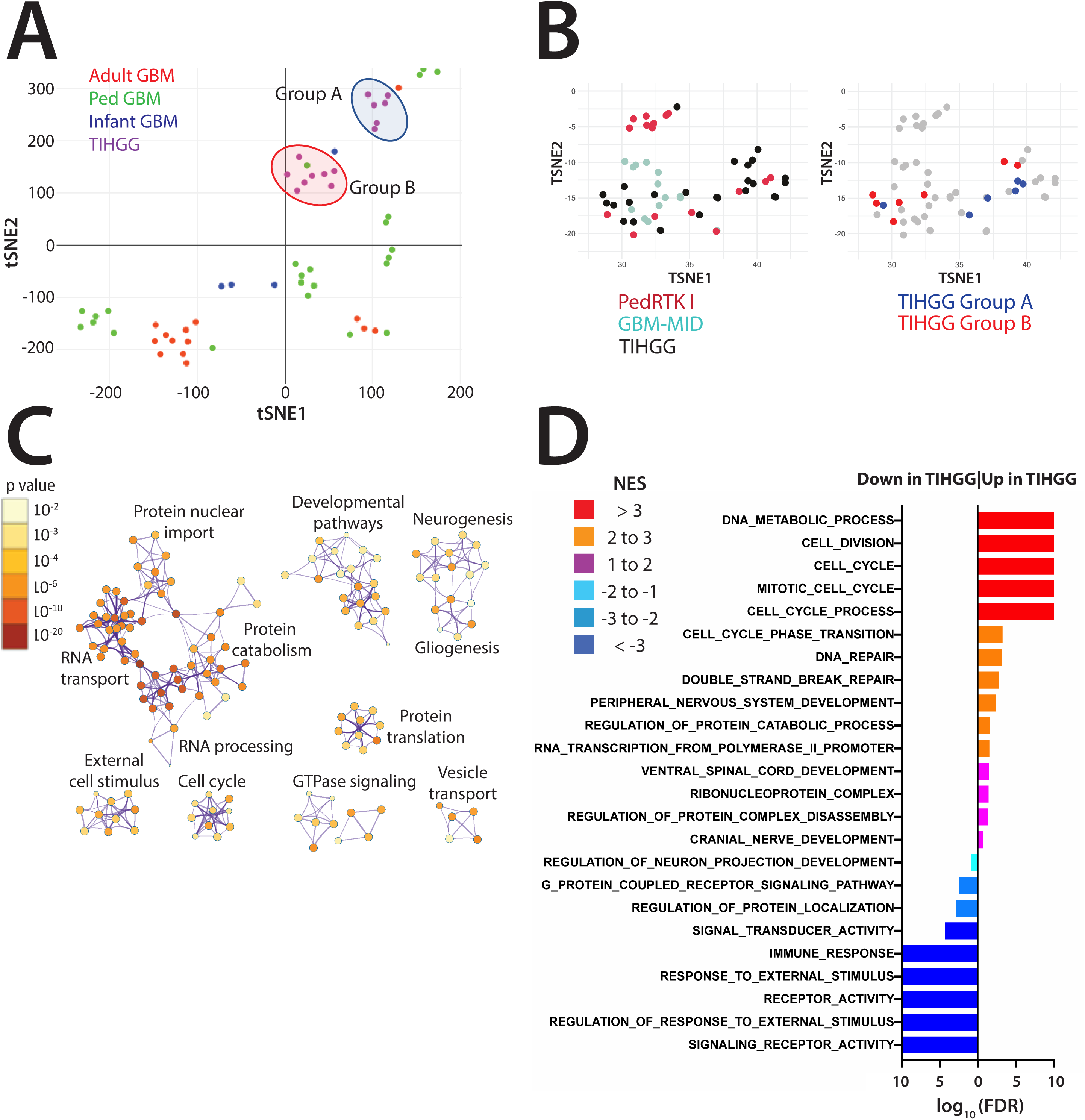
Gene expression profiling of TIHGG versus *de novo* GBM. A) Clustering of TIHGGs with microarray-based transcriptomic data (N=14) versus *de novo* pediatric GBM (N=24), infant GBM (N=4) and adult GBM (N=14) using t-SNE analysis. B) Methylation clustering showing locations of TIHGGs included in panel A; all TIHGGs included in panel A cluster in pedRTK I; methylation and transcriptomic clusters overlap with the exception of two samples. C) Cytoscape analysis based on gene ontology (GO) genesets show cellular pathways and processes that differ based upon gene expression between TIHGG and *de novo* GBM (from panel A); color scale represents p-value, and the comparisons are non-directional between sample sets. D) geneset enrichment analysis (GSEA) using GO genesets identifies differences in gene expression between TIHGG and *de novo* GBM (from panel A); number scale represents log of the false discovery rate (with FDR = 0 set to a value of 10 for purposes of plotting); colors represent the ratio of GSEA normalized enrichment score (NES) of TIHGG/*de novo* GBM.

To further investigate gene expression in TIHGG, we performed Metascape/Cytoscape analysis, which revealed several differences in gene ontology between TIHGG and the transcriptional reference group of *de novo* GBM. The differences include fundamental cellular processes, such as RNA processing and transport, protein translation and catabolism, and cellular signaling, as well as pathways controlling neurogenesis and gliogenesis (Fig. 6C, Sup. Table 16). To determine the directionality of gene expression patterns identified in the Metascape/Cytoscape analysis, we performed gene set enrichment analysis (GSEA) using the gene ontology (GO) gene set collection^17, 18^. The results of this analysis showed similarities in differential gene expression to that identified in the Metascape/Cytoscape analysis, with TIHGG samples showing relative enrichment of genes involved in DNA metabolism, cell cycle progression, DNA repair, nervous system development, and protein catabolism compared to *de novo* GBM, and relative depletion of genes related to immune response, signaling and cellular response to external stimulus, receptor activity, and neurogenesis (Fig. 6D, Sup. Table 17).

### Expression-based characterization of TIHGG subgroups

To characterize the TIHGG gene expression subgroups, we analyzed the differences in their relative gene expression, mutational status, and copy-number alterations. GSEA revealed that TIHGG gene expression Group A shows stem cell (proneural) and neuronal characteristics and has enriched expression of *MYC* pathway genes (Fig. 7A). Expression Group B has mesenchymal and astroglial characteristics, enriched expression of inflammatory (particularly NF-κB pathway) and immune genes, and depletion of DNA-repair pathway genes (Fig. 7A, Sup. Table 18). We did not observe relative expression differences between the TIHGG subgroups in cell cycle or proliferative genes, consistent with clinical experience showing that TIHGGs, in general, are highly proliferative.

**Figure 7.**
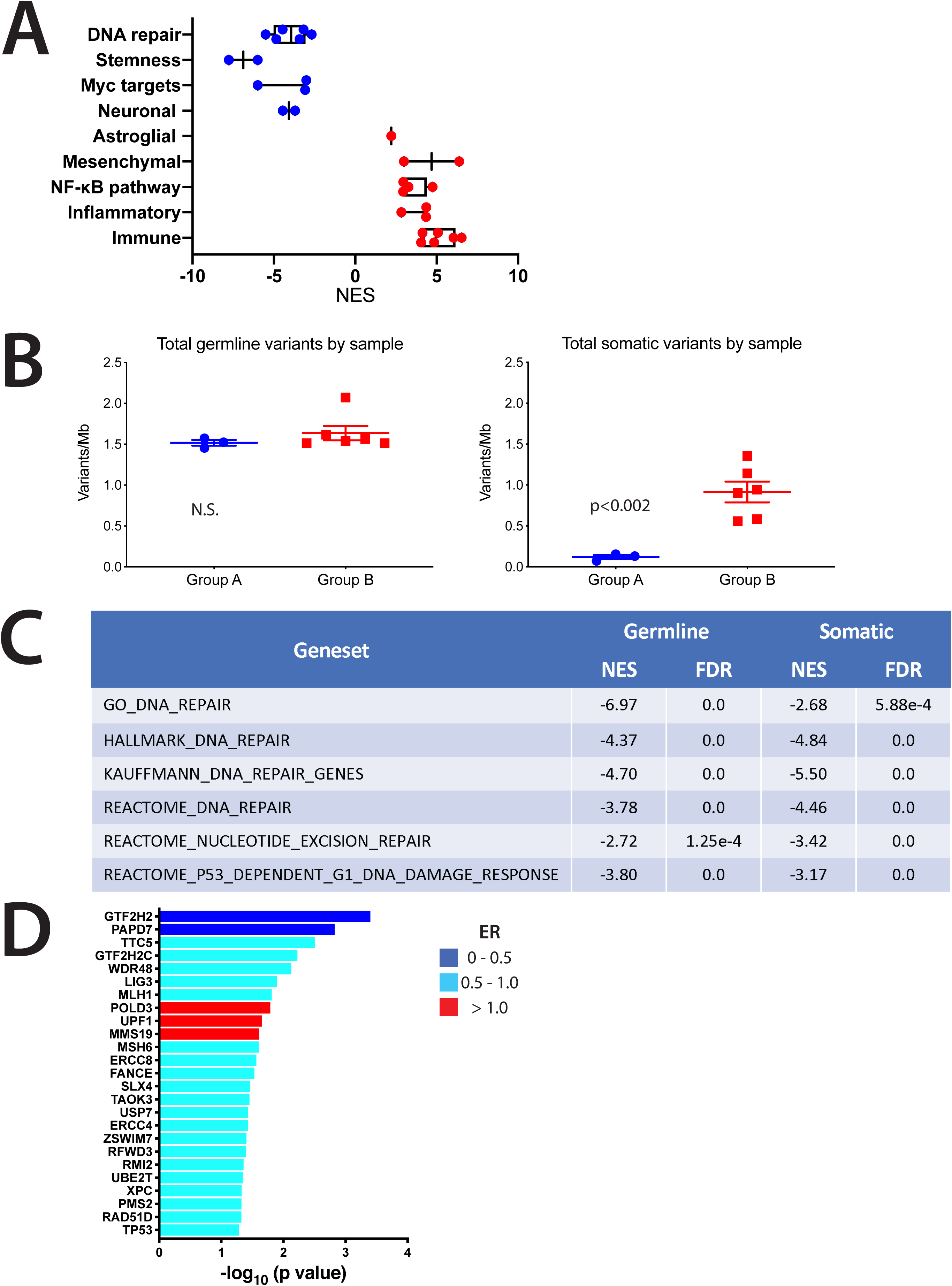
Gene expression and genetic profiling of Group A versus Group B TIHGG. A) GSEA results showing GO differences organized by category between Group A (blue) and Group B (red) TIHGG tumors; horizontal axis is NES for the comparison of TIHGG Group B vs. TIHGG Group A (NES > 0 means enrichment in Group B and NES < 0 means enrichment in Group A) (FDR values for comparisons are listed in Sup. Table 18); mean and SEM shown. B) Frequency of 1p loss and 1q gain based upon methylation profiling in TIHGG gene expression groups A and B; note that one TIHGG tumor for which there was transcriptomic data lacked a DNA sample and was not included in the methylation profiling; mean and SEM shown. C) For TIHGG samples with germline somatic genome sequencing data and transcriptomic data, germline variant load was identical between subgroups, but somatic load was approximately 9-fold greater in Group B (p<0.002); TIHGG sample with *MSH2* mismatch repair defect was excluded from this analysis. D) GSEA results for DNA repair pathways for Group B versus Group A germline (based on blood) and tumor samples; negative numbers represent depletion in Group B; Individual genes from the GO_DNA_REPAIR geneset with the most significant differences in expression between Group B and Group A are shown by decreasing p-value (top to bottom); colors represent the ratio of mean expression (ER) of Group B/Group A.

Using the DNA methylation data, we analyzed whether there were differences in copy-number alterations between the two gene expression subgroups. We found Group A to be enriched in 1p loss/1q gain (5/6 tumor samples) compared to Group B (0/7 tumor samples, p=0.005). GSEA showed particular differences at 1p34 (NES = −5.21, FDR=0) and 1p36 (NES = −7.77, FDR=0). The remaining gene- or chromosome-level amplifications and deletions identified in the TIHGG cohort, including Ch.13 or 14 loss, *PDGFRA* amplification, and *CDKN2A* loss, were relatively evenly distributed between the two TIHGG subgroups (Sup. Table 5). We did not identify patterns of significant point mutation or similar small-scale genetic differences between Group A and B tumors, nor did we find any relationship between initial malignancy and TIHGG subgroup (Fig. 5).

The relative depletion of DNA repair pathway genes that we identified in Group B relative to Group A led us to investigate whether there were differences in mutation load between the two groups. Excluding one hypermutator case, WGS data were available on matched blood and tumor samples for 3 cases from Group A and 6 from Group B. We found nearly identical levels of germline mutation load in Groups A and B (Fig. 7B). We found that Group B tumors, however, had a nine-fold greater somatic mutation load than Group A tumors (1.08 vs. 0.12 mut/Mb, p<0.002) (Fig. 7B).

As noted previously, relative to Group A, Group B is depleted in DNA repair pathway gene expression (mean NES = −3.96, p=0.001 vs. expected value of 1), which led us to compare DNA repair pathway gene expression in group A and B germline samples. We found that germline samples from Group A patients were likewise enriched in DNA repair pathway gene expression (mean NES = -4.30, p=0.001) (Fig. 7C). Individual DNA repair pathway genes with lower expression in Group B relative to Group A include *GTF2H2,* a subunit of RNA polymerase II transcription initiation factor IIH that is involved in nucleotide excision repair; *PAPD7,* a DNA polymerase involved in DNA repair; *ERCC4,* which is essential for nucleotide excision repair; and *TP53* (Fig. 7D, Sup. Table 19.). In a pairwise comparison of the individual DNA repair pathway genes shown in Figure 7D, the mean expression ratio (Group B/Group A) was 0.76 (p=9.79×10^-11^ vs. expected value of 1) (Sup. Table 19).

In summary, TIHGGs split into two transcriptional subgroups with distinct gene expression profiles relative to *de novo* GBM. Subgroup A resembles proneural GBM and is enriched in expression of *MYC*-pathway genes, while subgroup B resembles mesenchymal GBM and has greater mutational burden and decreased DNA repair gene expression.

### *In silico* and *in vitro* drug screening identify potentially effective therapeutic agents for the treatment of TIHGG

In order to investigate therapeutic susceptibilities in TIHGG tumors, we used *in silico* modeling to identify potential TIHGG sensitivity to drug classes compared to a set of 23 *de novo* pediatric and 14 *de novo* adult GBMs. We also performed an *in vitro* drug screen of two primary tumor-derived cell lines, one from gene expression Group A (MAF-496) and one from Group B (MAF-145), for sensitivity to a panel of all FDA-approved anti-cancer agents. The *in silico* modeling predicted several classes of therapeutic agents to have greater activity in TIHGG when compared with *de novo* GBM. These included DNA-damaging agents, microtubule polymerization/de-polymerization inhibitors, proteasome inhibitors, and histone deacetylase (HDAC) inhibitors (Fig. 8A, Sup. Table 20).

**Figure 8.**
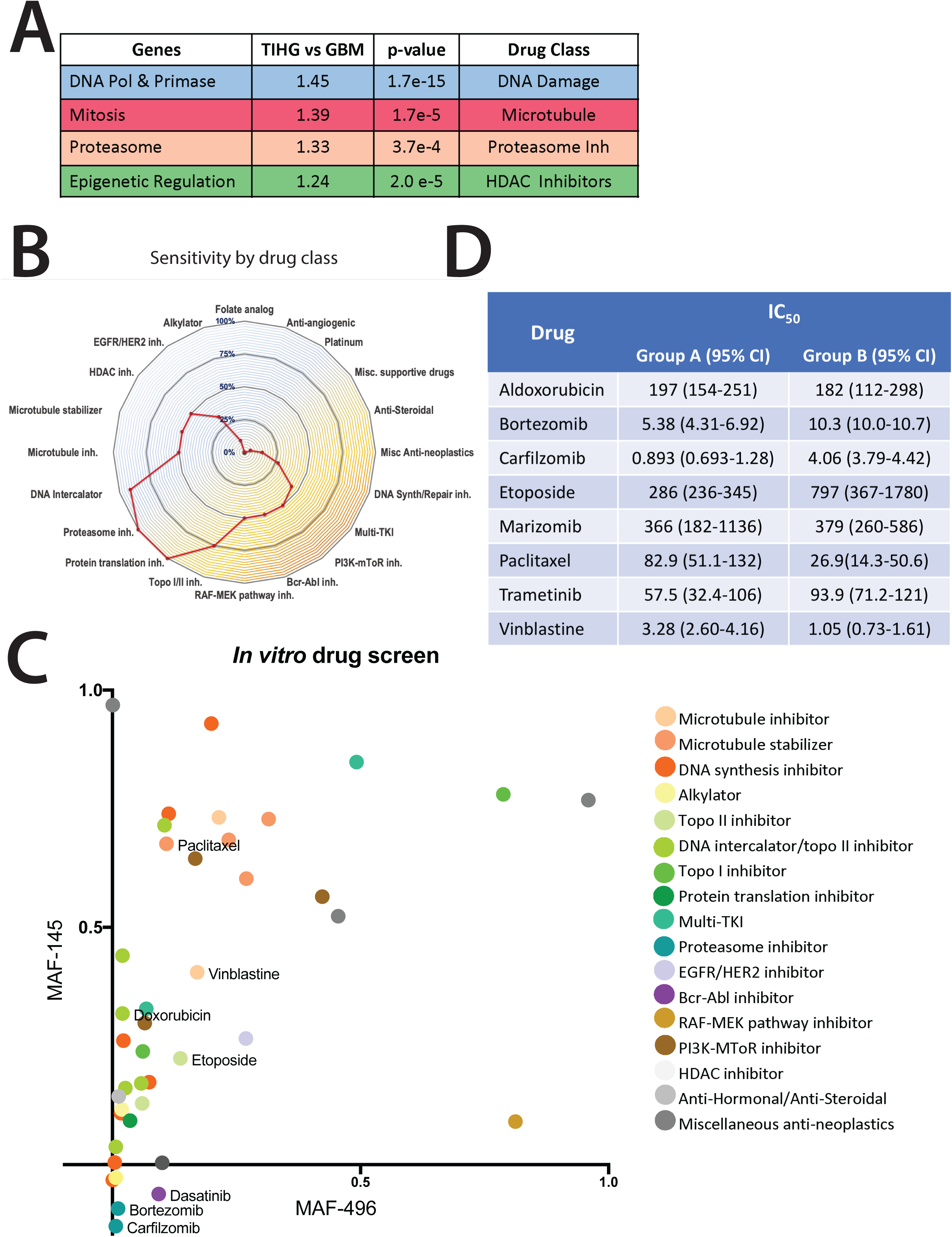
TIHGG preclinical drug screen. A) *In silico-*predicted response of TIHGG vs. *de novo* GBM tumors to drug classes based upon gene expression data. B) *In vitro* drug screening results by drug class for TIHGG cell lines MAF-145 (gene expression Group B) and MAF-496 (gene expression Group A) showing reduced survival by drug class relative to mean survival; screen performed using FDA-approved anti-cancer agents at 1 μM concentration. C) *In vitro* drug screen results in TIHGG cell lines using 1 μM concentration; Group A (MAF-496) cell line response is plotted on the x-axis and Group B cell line (MAF-145) on the y-axis. Dot colors correspond to drug class as shown in the legend. D) *In vitro* validation results in TIHGG cell lines MAF-145 and MAF-496 for candidate drugs identified through *in vitro* drug screen (panels B, D)

Our *in vitro* screen encompassed all FDA-approved anti-cancer agents, including multiple agents from the classes identified in the *in silico* screen, all dosed at 1 μM concentration (Sup. Figure 10A, Sup. Table 21). We found, combining results from the two cell lines, that 11/12 DNA intercalator results, 6/12 microtubule agent results, 4/4 proteasome inhibitor results, and 3/6 HDAC inhibitor results showed at least 50% cell death compared to vehicle (Fig. 8B, Sup. Table 21). The screen also identified topoisomerase I/II inhibitors (6/8 results) and PI3K-MTOR, BCR-ABL, and RAF-MEK pathway inhibitors (12/24 results total) as potentially effective using the 50% cell death criterion (Fig. 8B, Sup. Table 21). Folate analogs (0/6 results with at least 50% cell death), platinum drugs (0/6 results), and alkylators (3/32 results) performed poorly in the *in vitro* screen (Fig. 8B, Sup. Table 21). We found that, with the exception of proteasome inhibitors, which are further discussed below, the Group A cell line (MAF-496) was more susceptible to the effective drugs than the Group B cell line (MAF-145) (Fig. 8C).

To validate the screening results, we performed single-drug assays of selected agents at a range of concentrations in the two TIHGG cell lines. Drugs were selected for validation based upon the screening results and included two DNA-damaging agents (aldoxorubicin and etoposide), two microtubule agents (paclitaxel and vinblastine), three proteasome inhibitors (bortezomib, carfilzomib, and marizomib), and two kinase inhibitors (sunitinib, a multi-TKI inhibitor, and trametinib, a MEK inhibitor) (Fig. 8D, Sup. Fig. 10B-G). In the validation testing, aldoxorubicin (an anthracycline drug capable of blood-brain barrier penetration), etoposide, paclitaxel, and vinblastine showed sub-micromolar IC_50_ values in both cell lines (Fig. 8D, Sup. Fig. 10B-C), as did the MEK inhibitor trametinib (Fig. 8D, Sup. Fig. 10D). Sunitinib had an IC_50_ of 5,256 nM in the MAF-145 (Group B) cell line but did not reach an IC_50_ level in MAF-496 cells (Group A) (Sup. Fig. 10E). Validation testing of bortezomib and carfilzomib showed *in vitro* effectiveness in Group B cells, consistent with the enrichment of the NF-κB pathway in Group B tumors (Fig. 8D, Sup. Fig. 10F). In the Group A cell line, however, 25-45% of cells survived the highest concentration of each drug, suggesting the presence of a substantial drug-resistant population in Group A (Fig. 8D, Sup. Fig. 10F). We found that the proteasome inhibitor marizomib, which is capable of penetrating the blood-brain barrier, had IC_50_ values of 366 and 379 nM in the Group A and Group B lines, respectively, but also had a drug resistant population of approximately 30% of cells in both cell lines (Sup. Fig. 10G), which was not seen with bortezomib or carfilzomib. Further investigation showed that proteasome inhibition reduced nuclear levels of NF-κB, a known target of proteasome inhibition, in the Group B cell line (MAF-145) but had a lesser effect on already low nuclear levels of NF-κB in the Group A line (MAF-496) (Sup. Fig. 10H-I).

Taken together, our drug screening results in TIHGG tumors showed that drugs that interfere with S-phase (aldoxorubicin and etoposide) or M-phase (vinblastine and paclitaxel), and the MEK inhibitor trametinib, were effective *in vitro* in both TIHGG subtypes. In addition, proteasome inhibition was effective in the Group B cell line.

## Discussion

The present TIHGG cohort represents the largest and most comprehensive studied to date and extends prior knowledge from past studies that largely included targeted analyses of sequencing, expression, and staining of smaller tumor collections. The use of methylation profiling, WES/WGS of tumor and matched germline samples, RNA-seq, and drug screening help advance understanding of long-held questions regarding TIHGG, including similarities and differences from *de novo* pHGG, distinguishing TIHGG from recurrent tumors, TIHGG oncogenesis, subgroups within TIHGG, and potential therapeutic susceptibilities.

All but one of our cases had a clinical history of cranial radiation exposure. The exception was a high-risk neuroblastoma patient who received multiple rounds of multi-agent chemotherapy, including two cycles of high-dose chemotherapy, but only received radiation to her primary tumor site in the mediastinum, far removed from her frontal lobe TIHGG. Her latency and tumor findings from our assays cluster closely with the remainder of the cohort, suggesting chemotherapy was most likely causative. While it was exceptional that chemotherapy exposure alone should give rise to a TIHGG, the term treatment-induced is preferred to radiation-induced to reflect this possibility and the known interaction between combined modality therapy and increased incidence of subsequent malignancies^1^. In addition, since these tumors were a mix of WHO grade III, IV, and HGG NOS, we have replaced the term glioblastoma with high-grade glioma.

The results of our methylation and transcriptomic analyses delineate TIHGG as an entity separate from pHGG and identify two distinct transcriptional subgroups within the TIHGG cohort. Additional genomic and transcriptomic analyses identify characteristic genetic alterations and gene expression profiles that define the two subgroups. TIHGG were most frequently assigned to the pedRTK I methylation subgroup of pHGG, despite their varied clinical origin. Of note, resolving pedRTK I from IDHwt, GBM-mid subtype pHGG, and TIHGG was not possible in our analysis, thus highlighting the molecular and epigenetic similarities between these groups. One TIHGG case in our series clustered with diffuse midline glioma (DMG) but lacked the defining *H3K27M* mutation (Case 13). One recent case series described three patients who had received prior radiotherapy for medulloblastoma and later developed tumors clinically consistent with DMG^19^. While one case in our series had the H3K27M mutation (Case 7), this case lacked methylation data and was not be able to be clustered against reference cohorts. Because of the close association between the pedRTK I subgroup with TIHGG, specific consideration may be useful in future brain tumor taxonomy. The relative contribution of *de novo* HGG and TIHGG remains to be established. Additionally, whether exposure to ionizing radiation outside the therapeutic setting also predisposes to the tumor subclass is currently unclear.

We compared recurrent genetic alterations in TIHGG and *de novo* pHGG cases in the pedRTK I group to define potential differentiating characteristics. We observed an increased frequency of *CDKN2A* and *BCOR* losses in TIHGG. *BCOR* alterations are infrequently observed in pHGG without co-segregating *HIST1H3B* K27M mutations and are not described in pedRTK I pHGG^20^. While *BCOR* alterations noted in the cohort were predominately non-focal, the region showed a high frequency of overlap across TIHGG cases, suggesting potential biologic relevance. Of note, *BCOR* alterations noted in the Pediatric Cancer Genome Project were primarily due to frameshift or nonsense mutations and were often coupled with loss of heterozygosity^21^. Despite the increased frequency of *CDKN2A* deletions, we did not observe accompanying *BRAF* V600E alterations, a finding common in pediatric low-grade glioma and some pHGG^22^. Because we observed recurrent activation of MAP kinase pathway in our cohort, concomitant CDKN2A loss may be a necessary mechanism to prevent oncogene induced senescence, as has been described in other tumor types^23^.

The differentiation of TIHGG from recurrent pHGG or transformed low-grade glioma based on DNA methylation profiling remains challenging. Two cases in our cohort arose following focal radiotherapy for an optic pathway glioma but were localized to a region within the radiotherapy low dose region but distant from the original cancer and not coincident with a metastasis. Prior to review, these were expected to be transformation events, but after reviewing location of the TIHGG in each case relative to the original tumor and the radiotherapy field, it is clear that the TIHGG arose in previously uninvolved but irradiated normal brain. It remains unclear whether modern methods such as IMRT, which expose larger regions of normal tissue to non-therapeutic, albeit potentially mutagenic doses of radiation, will yield an increased incidence of TIHGG despite the increased conformality of the therapeutic dose region.

RNA-seq analysis differentiated the overall TIHGG cohort most distinctly from pHGG, which has been previously documented in array expression analysis^24^. Our results furthermore show distinct subgroups within the TIHGG cohort: a low mutation load group A that is enriched for Ch.1p loss/1q gain, has neuronal characteristics, and is enriched in expression of stemness genes and transcriptional targets of *MYC*, and a higher mutation load group B that is enriched in expression of inflammatory and immune pathway genes, has astroglial and mesenchymal characteristics, and is impaired in DNA repair processes. The subgroups do not correlate with original tumor type or histology. Instead, the depleted germline DNA repair pathway gene expression in the Group B patients suggests their potential susceptibility to develop TIHGG following mutagenic cancer treatment.

Given the potential for mutational profile to identify malignancy type and the difficulty of distinguishing recurrent primary from secondary tumors, we evaluated the frequency and type of somatic variants in TIHGG and pHGG. The relative frequency of pathogenic variants in all TIHGG in our cohort was not substantially increased relative to pHGG, as other groups have found^7^, but Group B TIHGG had increased mutational burden compared to Group A, suggesting heterogeneity in the pedRTK I/TIHGG subgroup. We also observed differential rates of select base changes in somatic non-coding regions. Increased frequency of variations in non-coding regions have also been observed in syndromes associated with genomic instability^25^, as mutations in coding regions are likely to result in negative selective pressure.

The therapeutic potential of photon radiotherapy in tumor tissue has always come at the cost of sub-lethal damage in adjacent normal tissue^26^. Irradiated normal tissue damaged directly by secondary electrons or indirectly by reactive oxygen species is known to produce non-lethal double-stranded DNA breaks, which lead to impaired function and survival in long-term survivors of childhood cancer^26^. Incomplete error-prone repair of genomic DNA via non-homologous end joining and non-error prone repair via homologous recombination result in genomic instability, which may be further modified by host normal tissue radiosensitivity, dictated by germline DNA damage response functionality.

TIHGG exhibits signs of a mutagenic origin, with oncogenic amplifications of Ch.1q, *PDGFRA* and *CDK4*, copy number losses in tumor suppressors including *CDKN2A* and *BCOR*, and pathogenic mutations in *TP53, NF1* and *MET* (fusion events). TIHGGs have more frequent *PDGFRA* gain and *CDKN2A* loss relative to *de novo* pHGG; TIHGG also displays increased expression of oncogenic pathways indicative of increased proliferation, including DNA metabolism and cell cycle genes. The characteristics of the two TIHGG gene expression subgroups suggest differences in tumorigenesis related to differing types of DNA damage and their respective DNA repair capabilities. Group A TIHGGs all have large-scale, chromosome-level abnormalities that are associated with poor outcome in pediatric brain tumors, including Ch. 1q gain and Ch. 13 loss. Gain of Ch. 1q distinguishes pediatric from adult HGG, has been recognized as a feature of radiation-induced tumors, and is an indicator of progression and poor outcome in ependymoma, neuroblastoma, and Wilms tumors^5, 27–30^. Specific associations between chromosome-level abnormalities and the other characteristics of group A tumors have not been reported but are an area for further inquiry. The higher mutation load of group B tumors could arise through the combination of mutagenic treatment-induced DNA damage and the downregulation of DNA repair pathways in these cases. Further insight into potential germline susceptibilities in group B patients related to impaired DNA repair may even lead to identification of patients at risk for TIHGG prior to primary tumor therapy and allow treatment modification to prevent TIHGG. Based on our germline data, we hypothesize these germline susceptibilities are different from known mutations causing tumor predisposition syndromes in that they may only manifest in the setting of highly mutagenic treatment.

Large-scale, localized DNA damage in the form of chromothripsis has also been recognized as a means of carcinogenesis following radiotherapy, as it can perpetuate a string of subsequent random molecular alterations. While pHGG exhibit genomic instability secondary to alterations in histone- and chromatin-modifying machinery, we observed an increased rate of chromothripsis in TIHGG relative to that of *de novo* pHGG. Chromothripsis was more common in the setting of pathogenic *TP53* mutations. This has also been observed in pediatric and adult malignancies not associated with therapeutic ionizing radiation^31^. Although chromothripsis is known to result in complex chromosomal alterations that can perpetuate a high frequency of copy-number alterations, we also observed characteristic broad and focal copy-number changes in non-chromothripsis cases (+1q, −6q, −13, −14, *+PDGFRA*, *+CDK4,* −*CDKN2A, −BCOR)* that have known oncogenic potential^27, 22, 32–34^. We observed several instances of oncogene amplification, likely facilitated by inclusion in eccDNA, which has been identified as a mechanism of resistance to targeted therapies. In tumor cells with eccDNA containing the oncogene *EGFR,* eccDNA levels increase in response to EGFR inhibition^35^. The eccDNA increase is frequently independent of and more efficient, in terms of onset and magnitude of response, than chromosomal copy-number alterations^35–37^. Similarly, serial samples from patients exposed to standard-of-care therapy show increases in *EGFR* levels during treatment with *EGFR* inhibitors, also likely in part facilitated by the presence of eccDNA^16^. These findings may explain the lack of observed efficacy of well-characterized, CNS-penetrant, targeted drugs such as PDGFRA inhibitors in our preclinical drug screens.

*In silico* and *in vitro* drug screening identified several FDA-approved drugs that merit further study as potential therapeutic agents to treat TIHGG. Drug classes identified as most effective against TIHGG (DNA-damaging agents, anti-mitotic drugs that target microtubules, and histone deacetylase (HDAC) inhibitors) have not been clinically effective in *de novo* pHGG, though clinical studies of HDAC inhibitors are ongoing.^38, 39^ Mechanistically, Group B TIHGG may show vulnerability to these DNA-damaging and anti-mitotic agents because of its DNA repair pathway deficiencies, as cells with substantial drug-induced DNA damage prevented from completing mitosis would be expected to undergo cell death associated with mitotic catastrophe.^40^ Interestingly, in the *in vitro* drug screen, the PARP2 inhibitor olaparib enhanced proliferation compared to vehicle in the Group B cell line (Sup. Table 21). Where DNA damage surveillance is intact, PARP2 inhibition is ineffective in inducing apoptosis and instead induces a choice of repair pathway, favoring error-prone, non-homologous end resection over homology-directed repair^41, 42^. Thus, in Group B TIHGG, in which homology-directed repair pathways are already deficient, PARP2 inhibition may enhance, rather than retard, proliferation by favoring non-homologous resection.

Proteasome inhibitors, which have not been studied beyond preclinical phases in pHGG, were effective *in vitro* in the Group B cell line, possibly through a mechanism involving inhibition of NF-κB-mediated inflammatory pathways. Clinically, proteasome inhibition as a therapeutic target is established in multiple myeloma and mantle cell lymphoma; however, its effectiveness is limited by acquired resistance^43^. This experience suggests that the efficacy of proteasome inhibition for TIHGG may depend upon identification of effective combination therapies. An interesting possibility for inhibiting the NF-κB pathway in conjunction with proteasome inhibition is through inhibition of MDA-9/Syntenin, which has shown promise in preclinical studies in reducing the invasiveness of radiation-induced GBM in adults.^44^ Targeting the MAPK pathway may also be an effective strategy given the frequent *NF1* mutations in TIHGG; MEK inhibition is a therapeutic strategy of interest in HGG and was effective (trametinib) in both TIHGG derived cell lines^45, 46^.

Strengths of the current study include the size and multi-institutional nature of the cohort, as well as the comprehensive, orthogonal, unbiased assays performed. Recent characterization of large cohorts of *de novo* HGG provide basis for comparing TIHGG to the HGG landscape in a way that has not previously been possible. We also characterized two primary human TIHGG cell lines, allowing for interrogation of therapeutic vulnerabilities. Weaknesses include the heterogeneity of the molecular assays performed across the cohort samples; this has been partially ameliorated through the performance of separate assays and analyses at only one institution per modality in order to standardize results. In addition, *in vivo* models of TIHGG are still in development, and should be the focus of future work. With cell lines available and common molecular alterations better understood, both patient-derived xenograft and genetically engineered mouse models are possible. Our study demonstrates similarities and differences between TIHGG and pHGG, as well as both homogeneity and heterogeneity within TIHGG. Therefore, more efficacious treatment regimens may include a common backbone incorporating some aspects of pHGG treatment, in combination with targeted treatments directed at specific molecular alterations and susceptibilities characterizing TIHGG subgroups and individual patient tumors.

## Methods

### TIHGG case review

Fifty-four cases were reviewed across multiple organizations (Children’s Hospital Colorado (CHCO), St. Jude Children’s Research Hospital, Childhood Cancer Survivors’ Study (CCSS), the University of Hamburg, and the University of Florida) from 1981 to 2015 to determine cohort eligibility. Initial query of the CCSS institutional tumor tissue bank was based on the history of prior radiation with subsequent development of HGG. Patients were enrolled on Institutional Review Board-approved protocols for the harvest and study of tissue for research, including Colorado Multi-Institutional Review Board (COMIRB 95-500) and SJCRH IRB Number: Pro00007403; Mnemonic: XPD17-029; Reference Number: 001628. After clinical review, 7 cases were judged to be recurrent primary pHGG, two cases were recurrent primary ependymoma, one case was found to be a recurrent vs. malignant transformation of a juvenile pilocytic astrocytoma, and one case was found to be a recurrent glioneuronal tumor. Tissue was not available for 6 cases, and tissue was of insufficient quality or quantity in 2 cases (Sup. Fig. 11).

We used a modified version of Cahan’s criteria to determine eligibility for tumors induced by radiation^47^. All radiation-induced tumors arose within the initial irradiated field. Although Cahan’s criteria specify that TIHGG must have a histologically proven difference between the initial and subsequent tumors, 7 cases had an initial diagnosis of glial origin (1 anaplastic astrocytoma, 2 astrocytomas, 3 ependymomas, and 1 ganglioglioma). The two cases with astrocytoma both presented as low-grade pilocytic astrocytomas arising in the optic tract. Both cases were treated with radiotherapy after progressive visual loss following a failed trial of chemotherapy. The resultant TIHGGs originated in the posterior fossa and occipital lobe, respectively, both within a region of normal but previously irradiated brain.

We also included tumors arising in two patients with germline predisposition syndromes in which clinical evidence supported the diagnosis of induction by radiation. One patient with known mismatch repair deficiency and germline *PMS2* mutation developed a primary anaplastic astrocytoma (AA) localized to the right frontal lobe at the age of 14 years and was managed with a gross total resection followed by erlotinib and adjuvant radiation as a part of SJHG04 (NCT00124657). Following 9.5 years of controlled disease, a new tumor was discovered centered in the left occipital lobe. The area was judged to be of sufficient latency to be designated a TIHGG. After review of the clinical history, radiographic imaging, initial radiotherapy plan, and pathology, it was felt that the subsequent GBM was more likely to be treatment induced, as it arose in the prior radiotherapy field, differed from the initial AA, arose from a region of normal brain parenchyma (except for previous irradiation), and occurred with significant delay, making a late recurrence of AA exceedingly unlikely. The second patient had a mismatch repair deficiency arising from a heterozygous loss of *MSH*. Upon presentation with Philadelphia chromosome-positive ALL at age 9 years, the patient received a bone marrow transplant and total body irradiation (12 Gy). At age 20 years, the patient developed a right frontal anaplastic oligodendroglioma (WHO grade III). The clinical diagnosis of a radiation-induced tumor was based upon the occurrence of the oligodendroglioma in the radiation field and the likelihood that radiation-induced DNA damage played a significant role in the tumor formation.

One tumor clinically diagnosed as being induced by prior exposure to chemotherapeutic agents was also included in the TIHGG cohort (Sup. Table 1)^47^. Details regarding the clinical history of all reviewed and included cases are in Sup. Fig. 1.

### TIHGG material available for analyses

Deidentified tumor tissue specimens from frozen or formalin-fixed paraffin embedded (FFPE) sections and frozen patient blood samples were processed following appropriate institutional review board approval (Fig. 1E). Whole-genome methylation analysis was completed in 31 cases. RNA was of sufficient quality for RNA-seq analysis in 14 cases. We conducted whole-genome sequencing on twelve frozen tumor and matched blood samples (to obtain germline genomic information) and whole-exome sequencing on 5 additional FFPE tumor samples^48^.

### Voxel-wise lesion mapping of TIHGG

T1 images were registered to the Montreal Neurological Institute template^49^ by using ANTsR^50^ and analyzed with voxel-based lesion symptom mapping (VLSM)^51^ to assess similarity between statistical maps by calculating the correlation between t-scores, treating lesion voxels as subjects.

### Patient-level statistical analysis

Patient- and sample-level statistical analyses were performed using Rstudio Version 1.1.463. Packages used for the presented analyses include “survminer”, “ggplot2”, “survival”, and “networkD3”. Continuous data are described using non-parametric measures of central tendency and are tested across strata using the Wilcoxon-Mann-Whitney test. Frequency data across groups were evaluated using either the Fisher’s exact test or Chi-square test. Time to event endpoints are summarized using the Kaplan Meier estimator. Differences in time to event strata are compared using the log-rank test.

### Methylation array processing

Tumor DNA was extracted from FFPE material by using the Maxwell16 FFPE Plus LEV purification kit and the Maxwell 16 instrument (Promega, Madison, WI) according to the manufacturer’s instructions. Extracted DNA from FFPE tissue underwent quality control assessment using the Illumina Infinium FFPE QC Assay kit for qPCR. The Delta Cq values for all samples were less than four. DNA concentration was assessed using PicoGreen. At least 300 ng of DNA was used per sample for the subsequent bisulfite conversion with the Zymo EZ-96 DNA Methylation kit. After the bisulfite conversion, the Infinium HD FFPE Restoration kit was used to restore degraded FFPE DNA to a state that is amplifiable by the Infinium HD FFPE methylation whole-genome amplification kit. Restored DNA was then plate-purified (with the Zymo ZR-96 DNA Clean & Concentrator-5), amplified, fragmented, precipitated, re-suspended, and hybridized to an Illumina Infinium Methylation EPIC 850K BeadChip array for 22 hours and 30 minutes (by using the Illumina Infinium Methylation EPIC assay kit). Following hybridization, the arrays were manually disassembled and washed. The subsequent X-Staining of the array features was processed on a Tecan Freedom EVO robotics system. The arrays were then manually coated and imaged using an Illumina iScan system with autoloader.

DNA methylation data analysis was performed using the open source statistical programming language R (R Core Team, 2016). Raw data files generated by the iScan array scanner were read and preprocessed using minfi Bioconductor package^52^. With the minfi package, the same preprocessing steps as in Illumina’s Genomestudio software were performed. In addition, the following filtering criteria were applied: removal of probes targeting the X and Y chromosomes, removal of probes containing - nucleotide polymorphism (dbSNP132 Common) within five base pairs of and including the targeted CpG-site, and removal of probes not mapping uniquely to the human reference genome (hg19), allowing for one mismatch. In total, 394,848 common probes of Illumina 450K and EPIC arrays were kept for clustering analysis.

### Statistical analysis of DNA methylation

To determine the subgroup affiliation of our TIHGG samples, we used the reference DNA methylation cohort published by Capper *et al*. and available from the gene expression omnibus (GSE90496)^12^. TIHGG samples were combined with the 2,801 reference IDATs containing CNS tumors and control brain tissues for unsupervised hierarchical clustering, as previously described^12^. In brief, the 32,000 most variable methylated CpG probes measured by standard deviation across combined samples were selected. Pearson correlation was calculated as the distance measured between samples, and the unsupervised hierarchical clustering was performed by the average linkage agglomeration method. The probe-level beta values were also analyzed using t-stochastic neighbor embedding (t-SNE)^53^. Hierarchical clustering and t-SNE analyses were repeated by using the top 12,000 differentially methylated probes against a reduced reference set of 666 tumors representing 16 different previously described methylation classes (Sup. Fig. 3, Sup. Table 7)^12^. The reduced set contained normal control methylation classes, high-grade diffuse astrocytic tumors, high-grade neuroepithelial tumor with MN1 alteration, pleomorphic xanthoastrocytoma, and anaplastic pilocytic astrocytoma. Supervised analysis was performed using the random forest DNA methylation class prediction algorithm (V11b2) by uploading raw IDAT files to www.molecularneuropathology.org^12^.

### Detection of copy-number alterations with methylation array data

Copy-number variation was analyzed with Illumina methylation arrays using the conumee Bioconductor package in R using default settings. The combined intensities of all available CpG probes were normalized against control samples from normal brain tissue using a linear regression approach. Platform-matched normal brain tissue controls were used to reference against EPIC 850k array or 450k array tumor samples. Mean segment value of −0.2 and 0.2 were used as thresholds for losses and gains, respectively. Copy-number plots were manually examined for selected copy-number alterations. When copy-number information was also available from sequencing data, the two results were compared and adjudicated. The adjudicated results are shown in Fig. 5 for selected recurrent somatic alterations.

### RNA-seq analysis

Library preparation was performed using the TruSeq Library Preparation Kit v2 (Agilent); directional mRNA sequencing was performed at the University of Colorado Anschutz Medical Campus Genomics and Microarray Core on a HiSeq 2500 sequencing system (Illumina) using single pass 125 bp reads (1×125) and approximately 50 million reads per sample. The resulting data were mapped to the human genome (hg19) by gSNAP, expression (FPKM) was derived by Cufflinks, and differential expression was analyzed with ANOVA in R. Output files contained read-depth data and FPKM expression levels for each sample and, where gene expression levels were compared between groups of samples, the ratio of expression in log_2_ format and a p-value for each gene. CICERO was used to detect fusion genes on the RNA-seq data^54^.

### Analysis of transcriptomic data

Using micro-array data (Affymetrix Human Genome U133 Plus 2.0 Array) previously acquired from tumor samples of patients treated at Children’s Hospital Colorado, we compared patterns of gene expression in 14 TIHGG samples (13 radiation-induced and one chemotherapy-induced) to those of a cohort of non-treatment-induced tumors consisting of 24 primary pHGG, 4 infant HGG, and 14 adult HGG. We performed clustering analysis using the t-SNE method available in the RTSNE package and confirmed the TIHGG subgrouping obtained through t-SNE by using non-negative matrix factorization (NMF)^53, 55^. Principal component analysis with 30 initial dimensions preceded the t-SNE analysis, in which we empirically selected perplexity of 3 and 50,000 iterations as providing optimal results. For the NMF analysis, we utilized the identical micro-array dataset used in the t-SNE analysis, using k (number of clusters) of 2–5. We performed Metascape analysis on the micro-array data using as input a list of 2,162 genes differentially expressed (p<0.01) between the TIHGG and HGG samples, followed by Cytoscape to identify differentially enriched pathways^56, 57^. We performed geneset enrichment analysis (GSEA, Broad Institute) to identify the direction of enrichment between the HGG and TIHGG groups and GSEA and Ingenuity Pathway Analysis (IPA, Qiagen) to identify gene expression patterns within the TIHGG cohort^58, 59^. GSEA results were evaluated using the normalized enrichment score (NES), in which increased expression results in a positive score, and reduced expression in a negative score. Scores were considered potentially informative from a statistical perspective if the false-discovery rate (FDR) was less than 0.25.

### Whole-genome sequencing

WGS library preparation and sequencing were performed by BGI Americas; 100-fold mean coverage data were acquired. BGI performed initial quality testing of the sample, including concentration and sample integrity/purity. Concentration was detected by fluorometer or microplate reader (*e.g.* Qubit Fluorometer, Invitrogen). Sample integrity and purity were detected by agarose gel electrophoresis (agarose gel concentration: 1%, voltage: 150 V, electrophoresis time: 40 min). Following quality confirmation, 1 μg genomic DNA was randomly fragmented by Covaris. The fragmented genomic DNA was selected by Agencourt AMPure XP-Medium kit to an average size of 200-400 bp. Fragments were end repaired and then 3’ adenylated. Adaptors were ligated to the ends of these 3’ adenylated fragments to facilitate amplification by PCR. PCR products were purified using the Agencourt AMPure XP-Medium kit. The double-stranded PCR products were heat denatured and circularized by the splint oligonucleotide sequence. The single-strand circular DNA (ssCir DNA) was formed as the final library qualified via a quality control procedure. The qualified libraries were sequenced by BGISEQ-500. Briefly, each ssCir DNA molecule was formed into a DNA nanoball (DNB) containing more than 300 copies through rolling-cycle replication. The DNBs were loaded into the patterned nanoarray using high density DNA nanochip technology. Finally, pair-end 100-bp reads were obtained by combinatorial Probe-Anchor Synthesis (cPAS). WGS mapping, quality control, and somatic mutation (SNV and INDEL) calling and classification have been described previously^60, 61^.^60, 61^. The tumor and germline reads were mapped to GRCh37-lite and Tier1 mutations (i.e. coding somatic mutations). CNA regions were identified using CONSERTING on the paired tumor-germline WGS samples with Log Ratio > 0.25 or < -0.25 reported^62^.^62^. Structural variants (SVs) of tumor WGS samples were identified using CREST based on soft-clipped reads evidence^63^.^63^. The SVs that have discordant reads support in tumor sample but not in paired germline sample were reported.

### Comparison of mutation frequency in de novo pHGG and TIHGG by expression group

We analyzed mutation load, small scale variants, and structural variants in the TIHGG tumor samples and matched blood samples. We noticed, based upon our initial review of the genome sequencing data, that several tumor samples appeared to have anomalously large numbers of mutations and decided to perform a mutation load analysis based upon that observation. We prepared vcf files containing QUAL-filtered (threshold of 50) and unfiltered putatively damaging variants. Germline mutation load was defined as the total number of mutations per sample in the QUAL filtered dataset.

Somatic mutation load was determined from tumor samples without QUAL-filtering. The mutation calls were filtered to require at least 4 reads per called variant in order to constitute a mutation. This approach was deemed reasonable because of the likely heterogeneity present in the tumor samples such that a clone constituting a fraction of the total sample could have an allelic-level mutation that would be detected in only a small fraction of the overall reads at a particular locus, resulting in a low QUAL score. Data were sensitivity tested to determine the thresholds for fold requirement and also to determine whether a QUAL threshold should be imposed. The tumor sample from the patient with Lynch syndrome was not included in this analysis because of the existence of a known DNA repair defect unrelated to the therapeutic radiation treatment.

### Whole-exome sequencing

Human genomic libraries were generated using the SureSelectXT kit specific for the Illumina HiSeq instrument (Catalog No. G9611B; Agilent Technologies, Santa Clara, CA), followed by exome enrichment using the SureSelectXT Human All Exon V6+COSMIC bait set (Catalog No. 5190-9307). The resulting exome-enriched libraries were then sequenced by the Genome Sequencing Facility on a HiSeq 4000 (Illumina). WES mapping and quality assessment have been described previously^61^.^60, 61^. The tumor reads were mapped to GRCh37-lite, and variants were called by Bambino^18^ and annotated by Medal Ceremony as “Gold”, “Silver”, “Bronze” or “unknown”^64^. “Gold” mutations and variants matching with COSMIC database were retained^65^. For other coding variants, those that were low-frequency (<0.001) or absent in ExAC/1000Genome/NHLBI databases were reported if the variant was supported with at least 5 mutant alleles and at least 30% VAF^66, 67^. The significance of mutated genes was assessed using the Significantly Mutated Gene test^68^. Mutation frequency and composition were analyzed by comparing the number and type of mutations across primary and TIHGG samples. The absolute number of mutations, and frequencies of base pair substitutions in SNVs were compared across TIHGG and primary HGG using a *t*-test and Chi-square test, respectively (Fig. 7C, Sup. Table 7, and Sup. Table 8). The frequencies of commonly altered genes in TIHGG and primary HGG were compared by using Fisher’s exact test and are listed in (Sup. Table 7).

### Evaluation of chromothripsis events and structure prediction of eccDNA

The presence or absence of chromothripsis was evaluated in twelve samples’ WGS data (Sup. Table 8). Four key criteria were used to infer chromothripsis as described by Korbel *et al*; (oscillating CNA regions, clustering of breakpoints, randomness of DNA fragment joins, and randomness of DNA fragment order)^69^. Chromothripsis was called when at least two criteria were satisfied and further evaluated by manual review. The eccDNA structures were constructed following the procedures described in Xu *et al.* by identifying the cyclic graphs composed of highly amplified CNA segments and their associated SVs^16^.

### FISH analysis

Dual-color FISH was performed on 4-µm thick paraffin-embedded tissue sections. Probes were derived from BAC clones (BACPAC Resources, Oakland, CA) and labeled with either AlexaFluor-488 or AlexaFluor-555 fluorochromes. BAC clones were used to construct probes for the following genes: *PDGFRA (*laboratory-developed probe (RP11-231C18 & 601I15); 4p control (CTD-2057N12 & CTD-2588A19), and *CDK4* (Empire Genomics, Williamsville, New York, Cat# CDK4-CHR12-20-ORGR). Probes were co-denatured with the target cells on a slide moat at 90°C for 12 minutes. The slides were incubated overnight at 37°C on a slide moat and then washed in 4M Urea/2xSSC at 25°C for 1 minute. Nuclei were counterstained with DAPI (200ng/mL) (Vector Labs) for viewing on an Olympus BX51 fluorescence microscope equipped with a 100-watt mercury lamp; FITC, rhodamine, and DAPI filters; 100X PlanApo (1.40) oil objective; and a Jai CV digital camera. Images were captured and processed using the Cytovision software from Leica Biosystems (Richmond, IL)^70^.

### In silico and in vitro drug screening and validations

In silico *screen*. Micro-array data for 12 TIHGG samples and 37 *de novo* GBM samples (23 pediatric, 14 adult) were compared to target genes of several drug classes. A ratio of mean expression and p-value for each target gene in the TIHGG and *de novo* GBM samples were computed, and an overall ratio and p-value of all target genes used for each therapeutic classification for each target gene were computed and reported (Supp. Table 20).

In vitro *drug screen and validation:* For the drug screen performed in TIHGG cell lines MAF-145 and MAF-496, we utilized the Approved Oncology Drugs Set VI (National Cancer Institute), comprising 129 drugs, supplemented by selinexor (Karyopharm Therapeutics) and AZD2014 (Astra Zeneca). The complete list of drugs included in the screen is included in Sup. Table 21. Cells were plated at a density of 5,000 cells per well in in 90 μL medium in a 96-well treated cell culture plate (Corning #3595) and allowed to gain adhesion overnight. Drugs were applied in 10 μL of medium/1% DMSO at a concentration of 10 μM, resulting in a final concentration of 1 μM and 0.1% DMSO. Cells were incubated in drug for 5 days. DMSO (0.1%) was utilized as a control. Cell viability was assayed following 5 days of treatment using incubation with tritiated thymidine and quantification using a scintillation counter. Results were collected as counts/min and converted to survival using the formula (sample – medum)/(DMSO – medium) where “sample” is the scintillation count for each drug-treated sample, “medium” is the scintillation count for a well containing medium only, and “DMSO” is the scintillation count for a three-well average of cells treated with 0.1% DMSO only (Sup. Table 21). The drug screen was conducted twice in MAF-145 cells and once in MAF-496 due to limitations on cell availability. Validation tests of single drugs were conducted using drug concentrations ranging from 0.316 nM to 10 μM in half-log_10_ increments. Cells were plated as described above and incubated in drug for 120 hours. Three biological replicates were used for each drug concentration. Results were assessed using CellTiter 96 Aqueous One Solution Cell Proliferation Assay (Promega), according to manufacturer instructions. Survival was computed as above. IC_50_ values were calculated using a variable slope four parameter non-linear model with maximum survival constrained at 100% (Prism 7, Graphpad).

*Immunofluorescence staining of in vitro samples:* Cells were plated at a density of 20,000 cells per well in BioCoat chamber slides coated with poly-D-lysine or poly-D-lysine and laminin (Corning) and allowed approximately 24-48 hours in which to develop adhesion before subjecting them to experimental conditions. Cells to be stained were fixed for 20 minutes in formaldehyde diluted to 3.7% in PBS (Sigma), permeabilized in 0.1% Triton-X in PBS for 10 minutes, and blocked for 45 minutes in 4% BSA in PBS supplemented with 0.05% Triton-X. Cells were incubated in primary antibody to the p65 subunit of NF-κB (Cell Signaling, #6956, 1:400) diluted with 4% BSA (in PBS and 0.05% Triton-X) for 1h at room temperature or overnight at 4**°**C. Following multiple rinses with PBS, cells were incubated in a secondary fluorophore (Alexa Fluor 488) for 1 hour, rinsed, and coverslips were then adhered using ProLong Gold antifade reagent with DAPI (Invitrogen). Confocal imaging was performed at 400x using 405 nM (DAPI) and 488 nM (Alexa Fluor 488) lasers on a 3I Marianas imaging system (Intelligent Imaging Innovations). Images were captured using an Evolve 16-bit EMCCD camera (Photometrics).

## Acknowledgements

We thank members of the Zhang and Wu laboratories (St. Jude Children’s Research Hospital) and members of the Foreman and Vibhakar laboratories (University of Colorado) for assistance and discussion. We also thank Matthew Lear (St. Jude Children’s Research Hospital) for his help in acquiring and performing preliminary clinical analysis of primary tumor samples and Scott Olsen, Granger Ridout, and Emily Walker (Hartwell Center for Bioinformatics and Biotechnology) for their assistance in sequencing samples. RNA-seq library preparation and high-throughput sequencing were conducted at the University of Colorado Cancer Center Genomics Shared Resource; microscopy was performed at the University of Colorado Denver Advanced Light Microscopy Core Facility; and tumor sample preparation for imaging was performed by the University of Colorado Cancer Center Histology Shared Resource. This work was supported by the National Cancer Institute (CA55727, G.T. Armstrong, Principal Investigator), the Fördergemeinschaft Kinderkrebs-Zentrum Hamburg (US), St. Jude Children’s Research Hospital Cancer Center Support (CORE) grant (CA21765, C. Roberts, Principal Investigator), ALSAC, the Morgan Adams Foundation (ALG), and a St. Baldrick’s Foundation Scholarship (ALG).

## Supplementary Figures

**Supplementary Figure 1.**
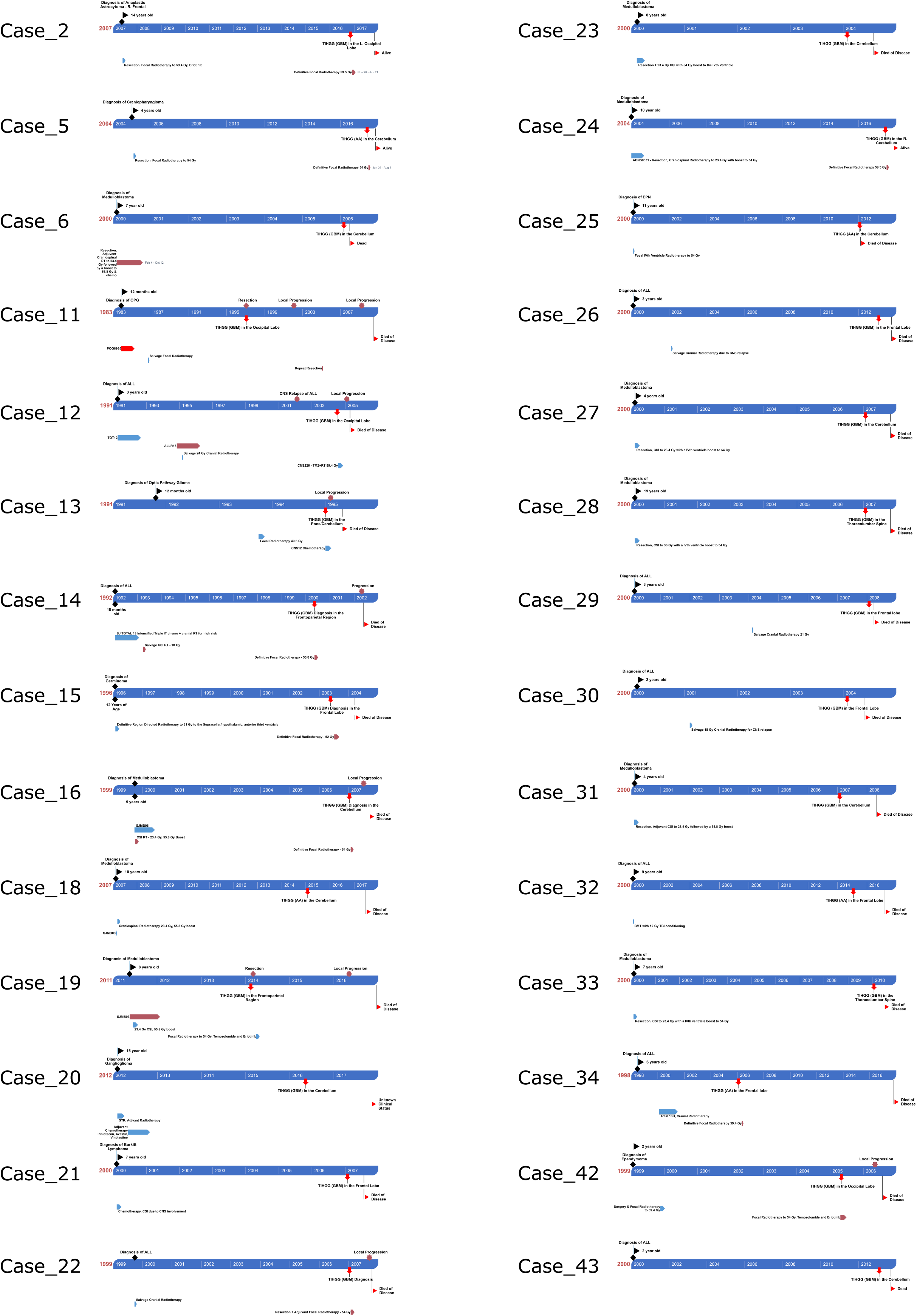
Case Treatment Histories.

**Supplementary Figure 2.**
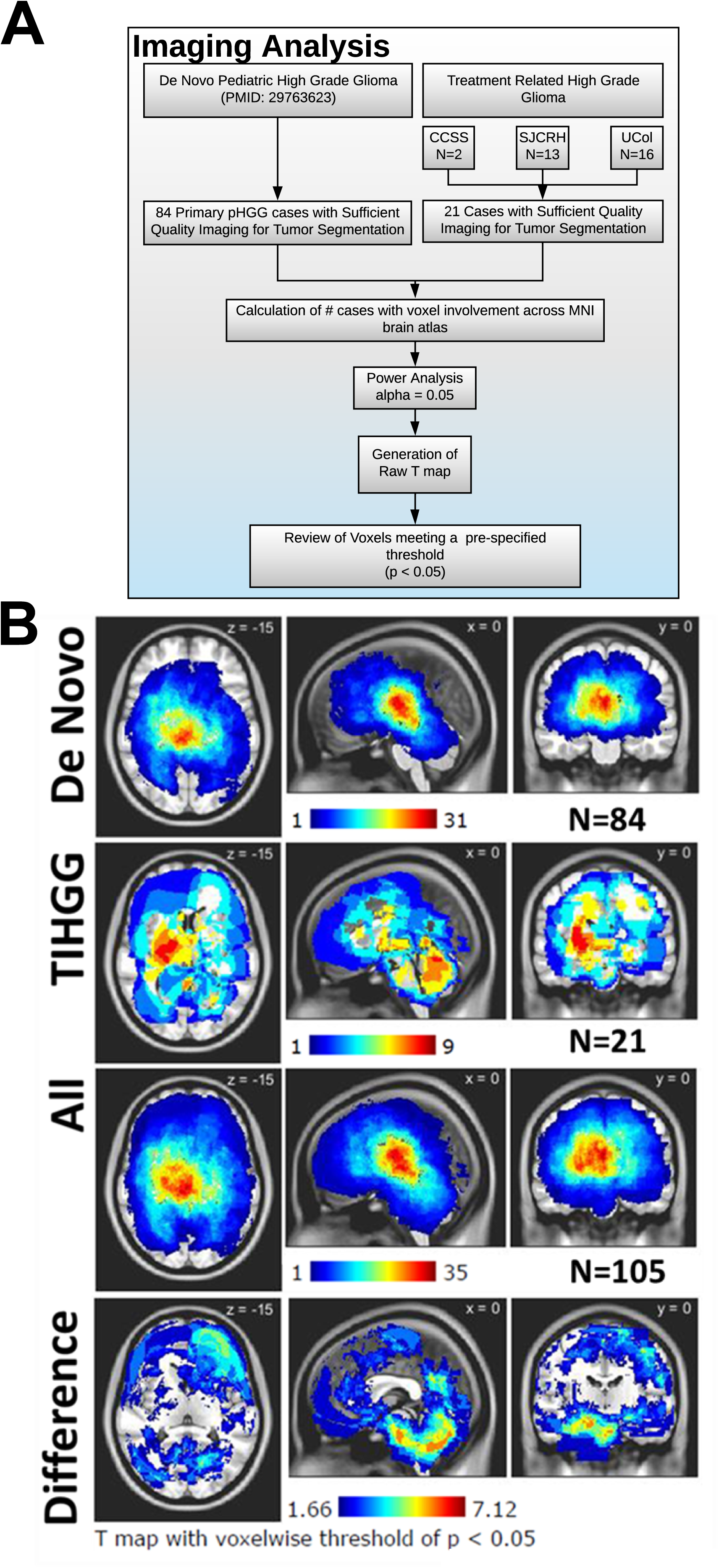
Anatomical distribution of *de novo* pHGG and TIHGG at initial presentation. A) (Top Row) Distribution of 88 “*de novo*” pHGGs; (2nd Row) 18 TIHGG; (3rd Row) Combined distribution of both *de novo* pHGG and TIHGGs; (Bottom Row) Spatial differences in distribution between *de novo* pHGGs and TIHGG using voxel-based lesion mapping (VLSM). B) Imaging analysis schema.

**Supplementary Figure 3.**
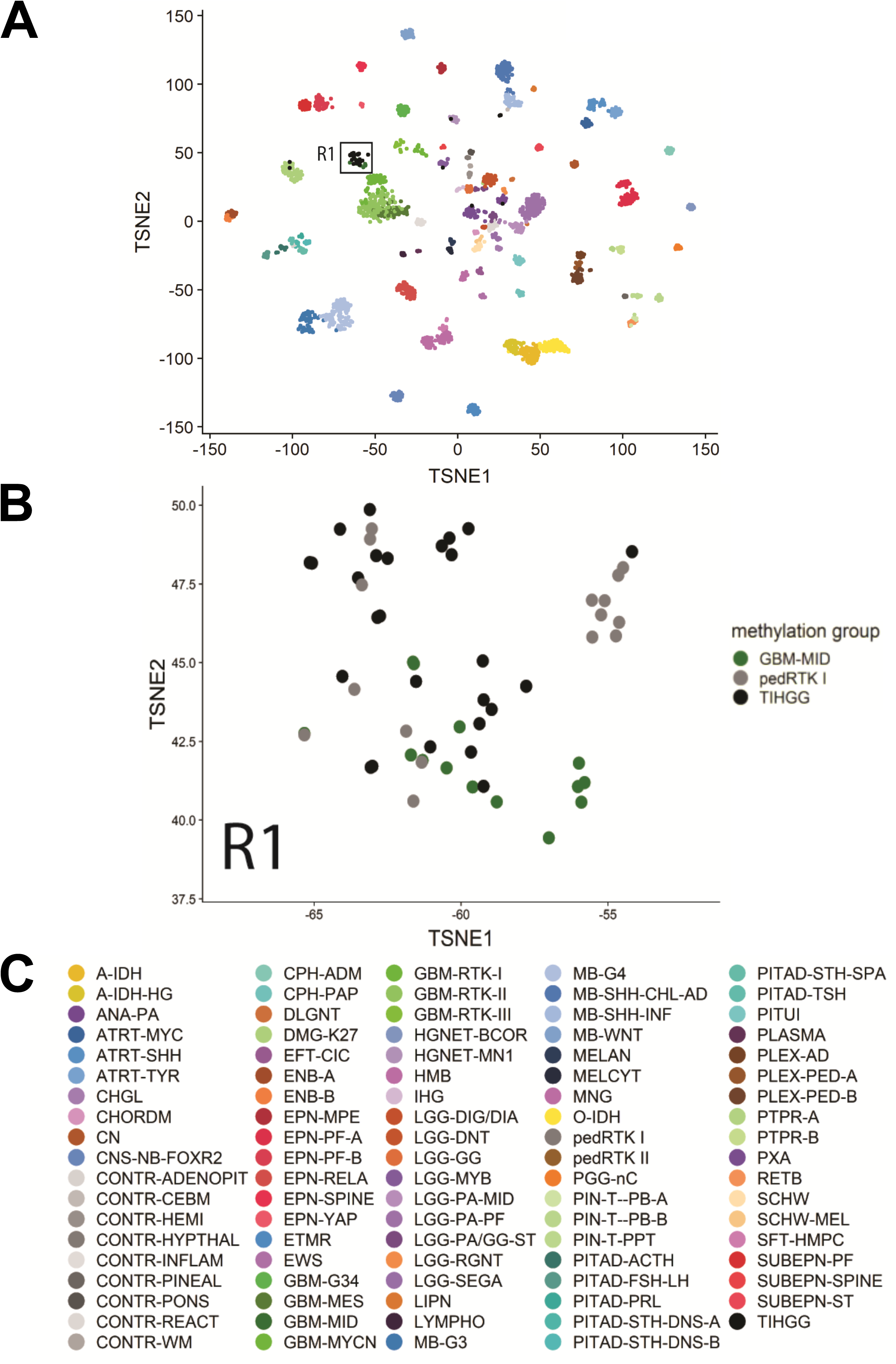
Position of TIHGG among *de novo* CNS tumors. A) Localization of TIHGG relative to other CNS cancers in t-SNE space. B) Magnified region indicated by A1 illustrating the close relationship between GBM-MID and TIHGG. C) Diagnosis key.

**Supplementary Figure 4.**
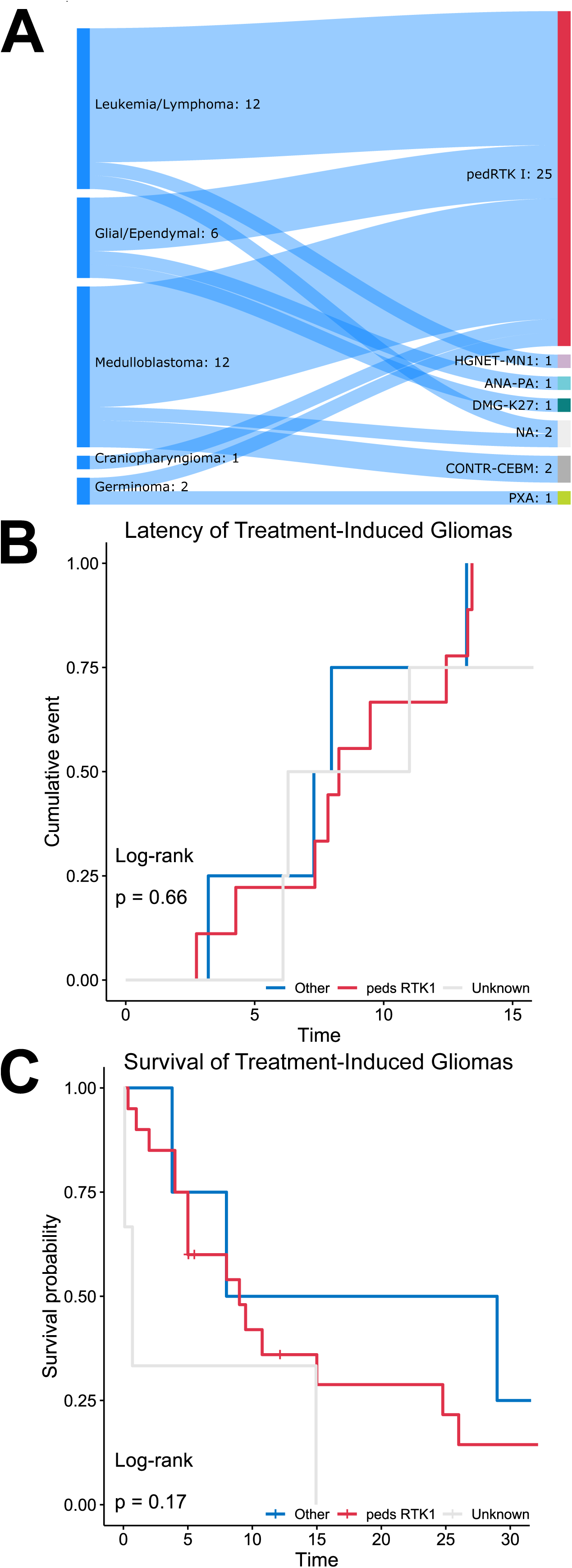
Results of clustering of TIHGG cases. A) Sankey diagram illustrating the inter-relationship between initial diagnosis and methylation-defined CNS tumor subgroup. B) Latency of TIHGG by consensus methylation cluster. C) Overall survival of TIHGG cases stratified by pedRTK I vs. other vs. unknown.

**Supplementary Figure 5.**
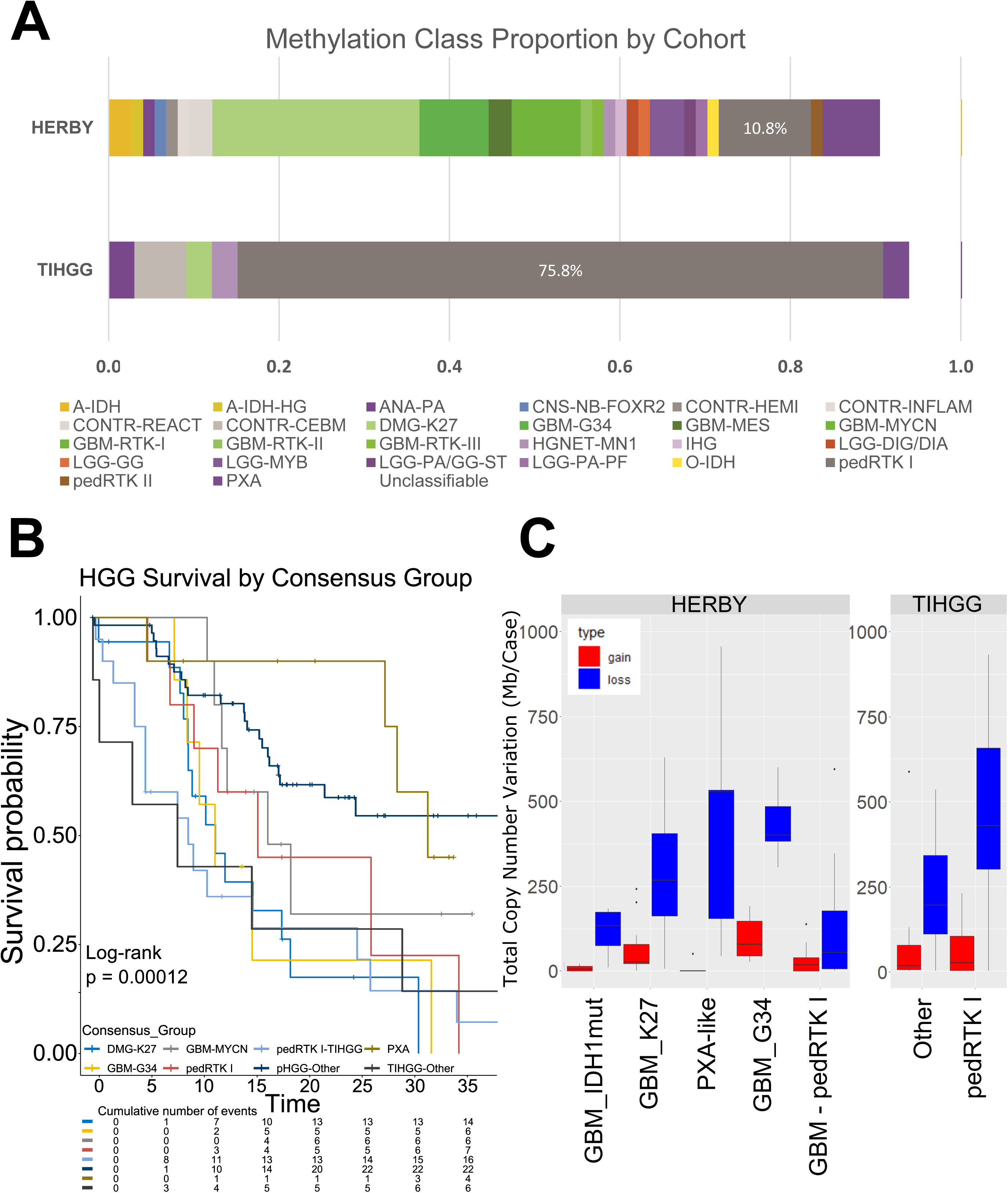
Comparison of TIHGG to *de novo* pHGG. A) Proportion of consensus cluster subtypes of TIHGG and primary pHGG cases from the HERBY dataset. B) Survival of TIHGG and pHGG stratified by consensus methylation group. C) Total genomic length (Mb) lost and gained by methylation subgroup; boxes show median and 1^st^/3^rd^ quartiles, with whiskers representing range limited to 1.5x the interquartile range from the box edge.

**Supplementary Figure 6.**
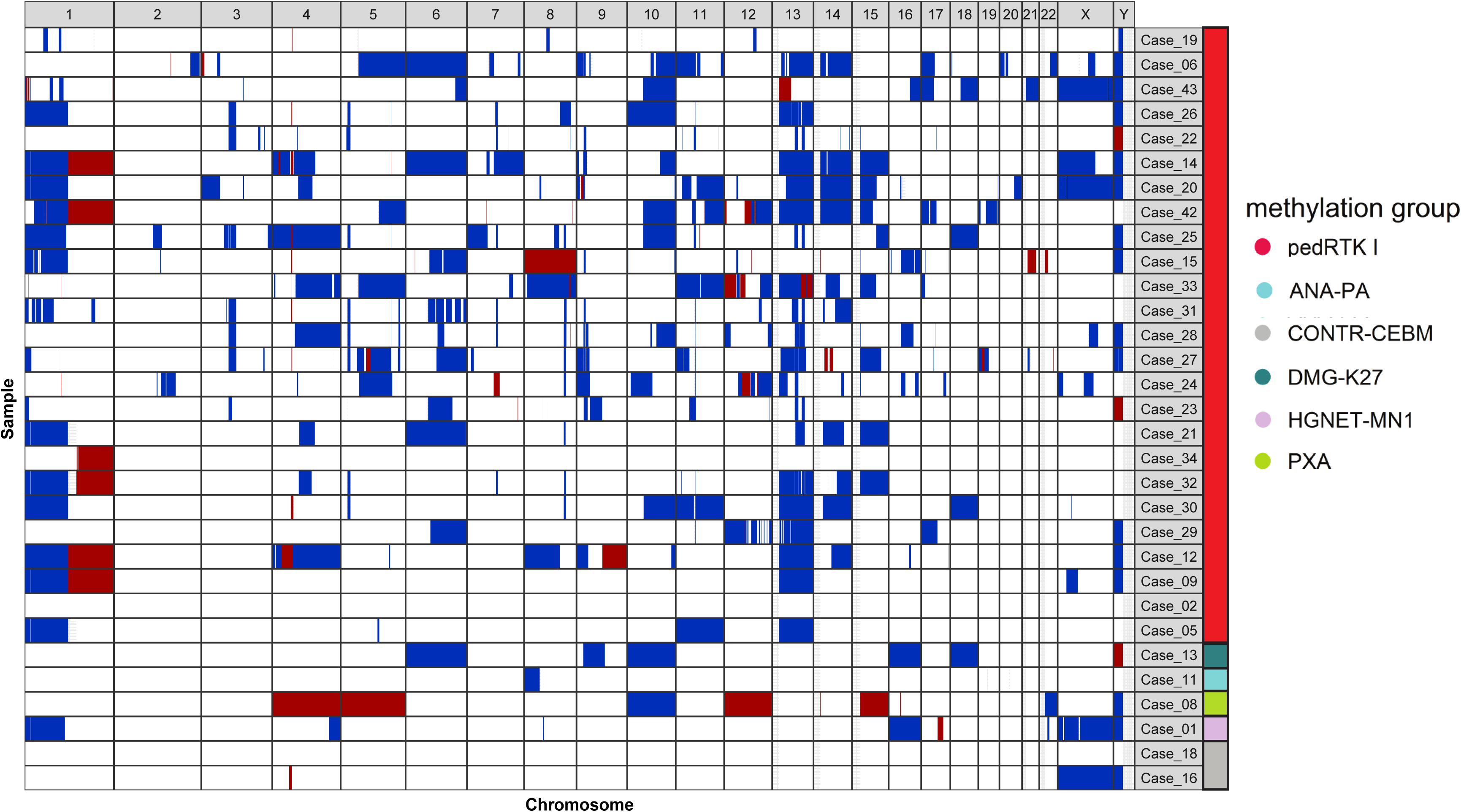
Spectral Plot illustrating chromosome-level copy-number alterations across TIHGG cases.

**Supplementary Figure 7.**
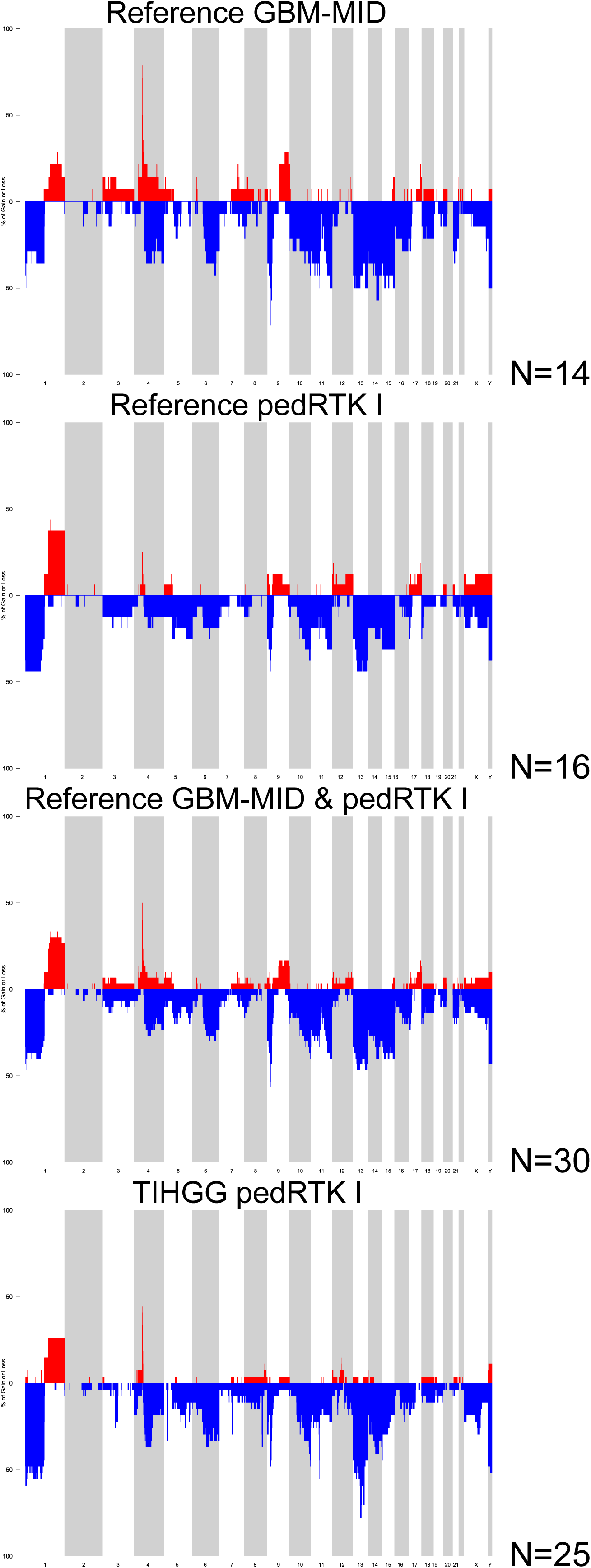
Comparison of copy-number alterations in TIHGG and *de novo* pHGG stratified by consensus methylation class.

**Supplementary Figure 8.**
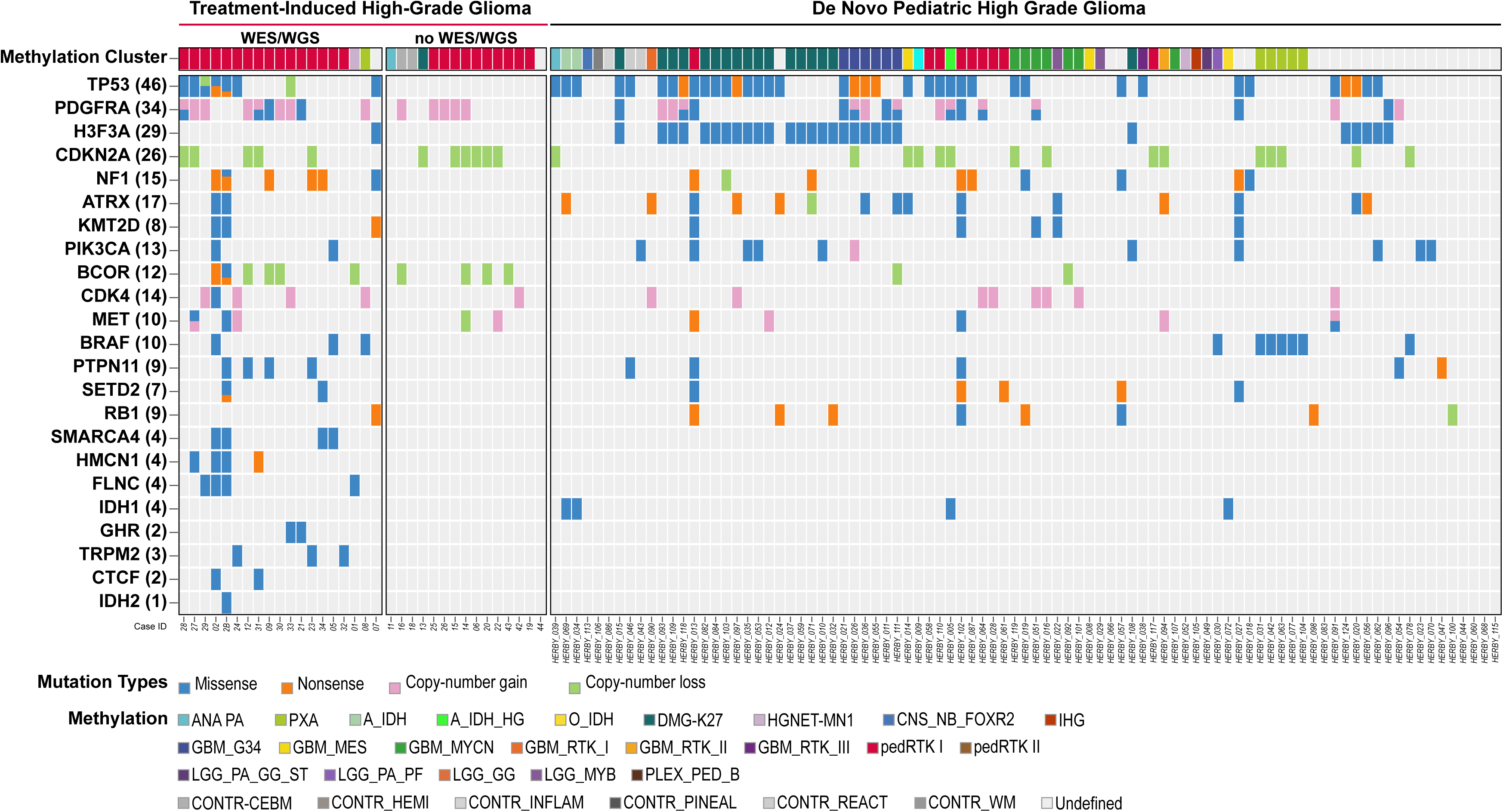
Recurrent molecular alterations in TIHGG relative to *de novo* pHGG (HERBY dataset). Oncoprint describing the clinical characteristics, histopathological features, methylation profile, tier1-mutations, genes affected by copy-number gain/loss, and fusion genes in TIHGG and *de novo* pHGG.

**Supplementary Figure 9.**
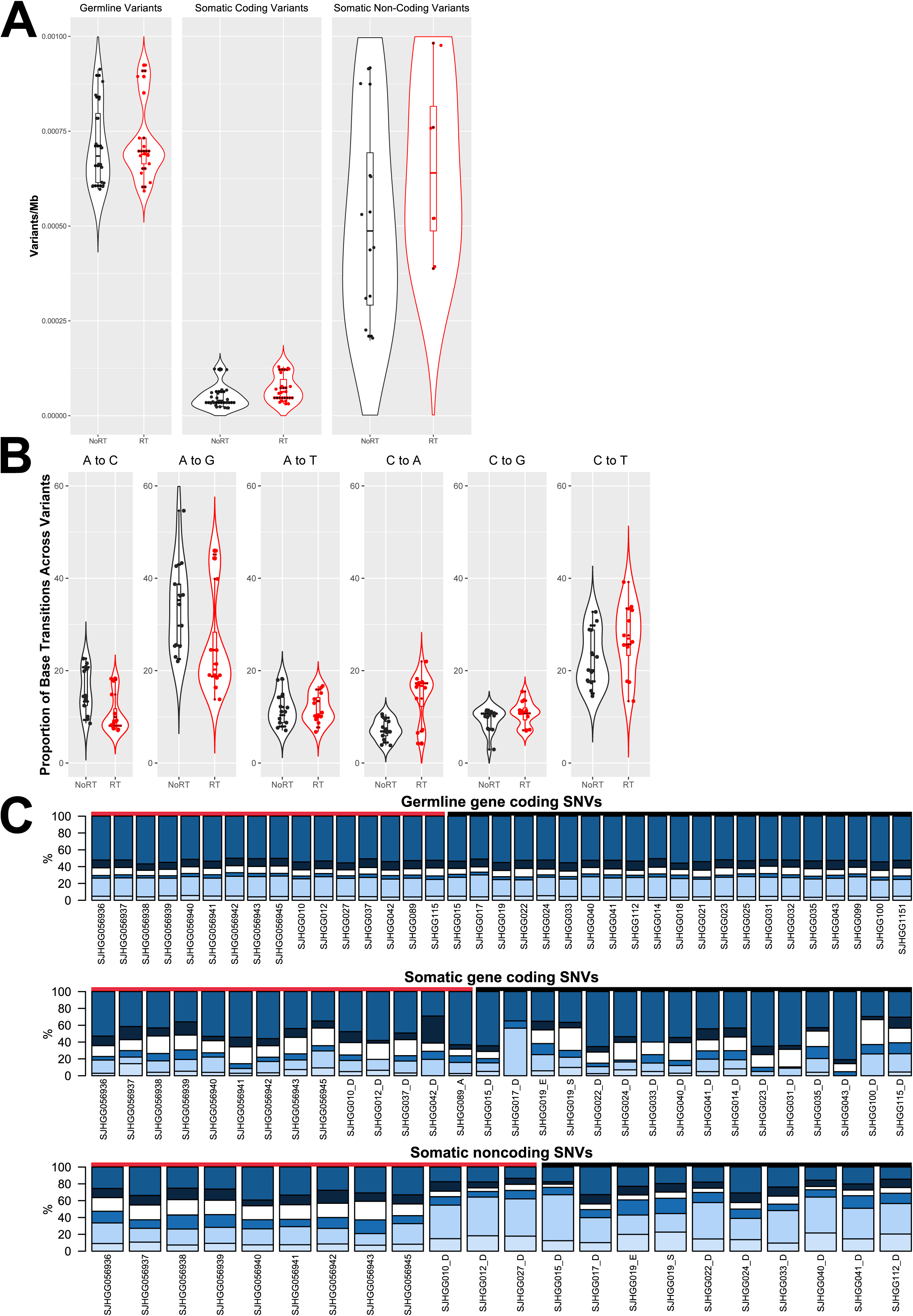
Germline and somatic variants in TIHGG relative to *de novo* pHGG (Wu G reference cohort) stratified by base transition type. A) Total number of germline, somatic coding, and somatic non-coding variants per Mb stratified by *de novo* pHGG vs. TIHGG. B) Proportion of base transitions in somatic non-coding variants per Mb stratified by primary pHGG vs. TIHGG. The relative frequency of base transitions in germline coding SNVs. C) Somatic coding SNVs and somatic non-noncoding SNVs. For violin plots, box shows median and interquartile range, whiskers show 95% confidence interval, total length of violin represents range, and width of violin shows frequency.

**Supplementary Figure 10.**
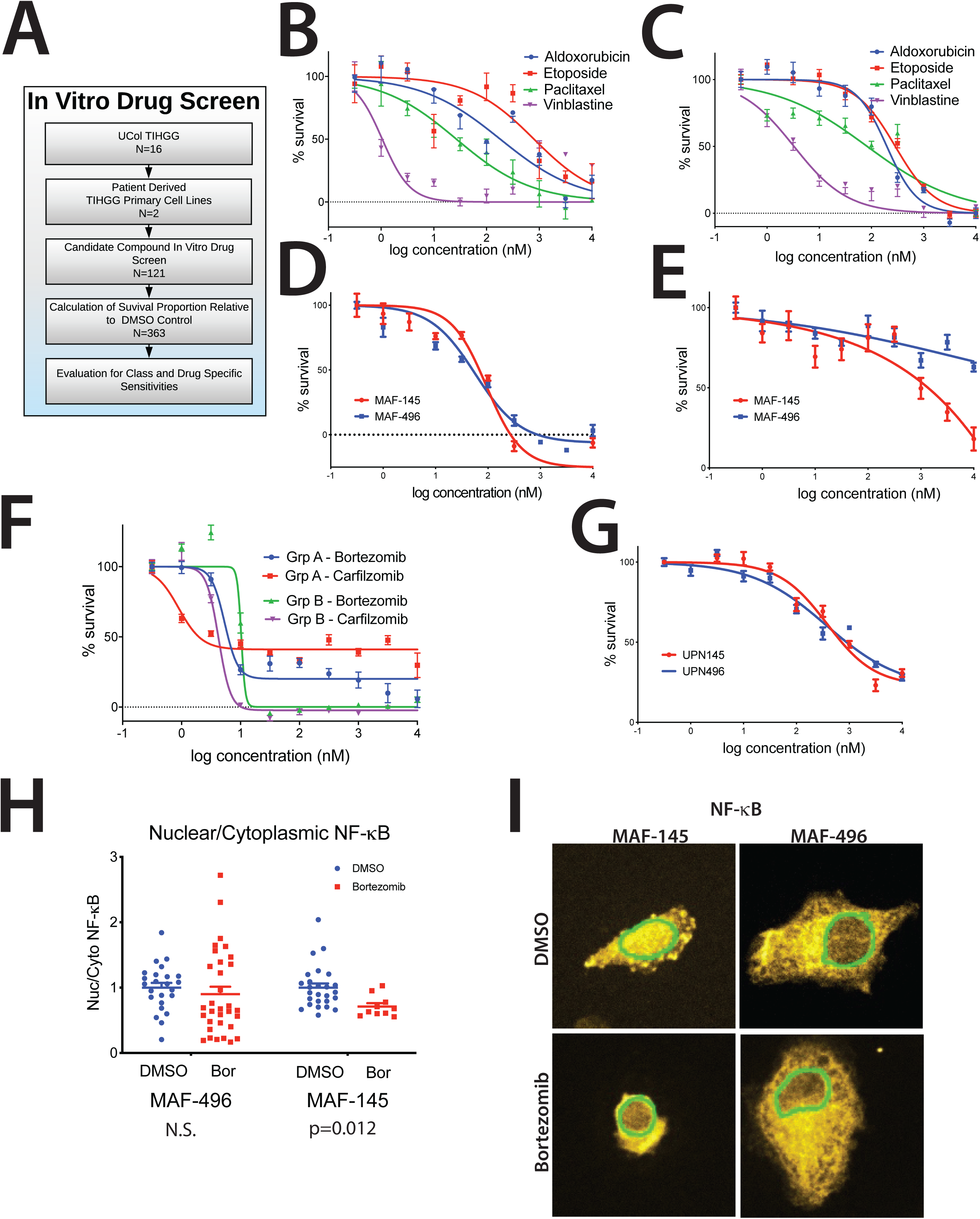
TIHGG *in vitro* sensitivities by expression profile. A) *In vitro* drug screen consort diagram; primary cell lines derived from TIHGG tumors were MAF-145 (gene expression TIHGG Group B) and MAF-496 (gene expression TIHGG Group A). B) MTS assay results for indicated drugs in MAF-496 (IC_50_ values listed in Fig. 8D). C) MTS results for indicated drugs in MAF-145 (IC_50_ values listed in Fig. 8D). D) MTS results for trametinib. E) MTS results for sunitinib. F) MTS assay results for proteasome inhibitors bortezomib and carfilzomib in MAF-145 and MAF-496 cell lines. G) MTS results for marizomib. H) NF-κB levels in Group A (MAF-496) and B (MAF-145) TIHGG cell lines treated with vehicle and bortezomib. I) Immunofluorescence of NF-κB nuclear localization in vehicle bortezomib-treated Group A (MAF-496) and Group B (MAF-145) TIHGG cell lines. Mean and SEM shown.

**Supplementary Figure 11.**
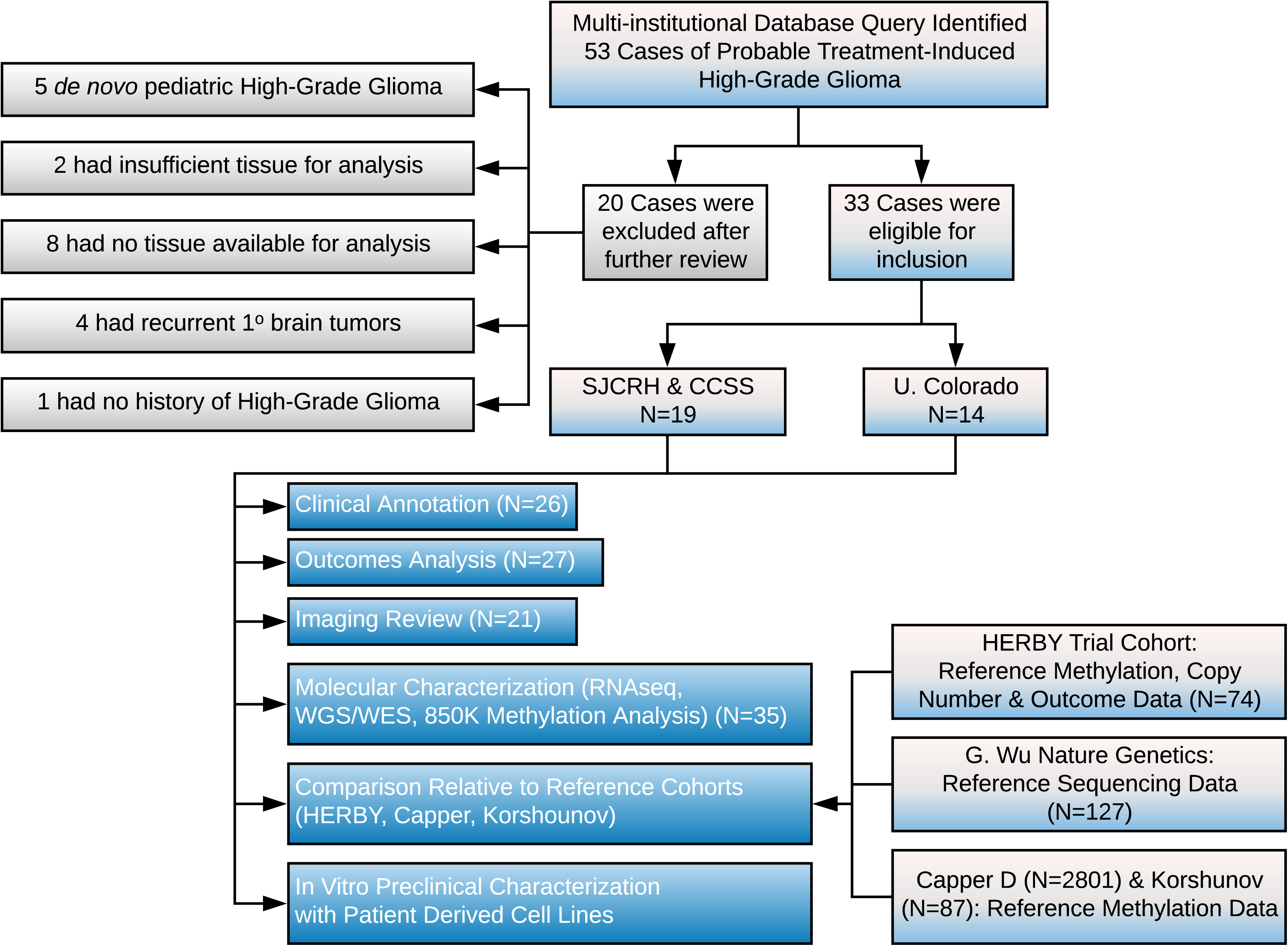
Study consort diagram. CCSS = Childhood Cancer Survivor Study, SJCRH = St. Jude Children’s Research Hospital, U. Colorado = University of Colorado, WGS = whole-genome sequencing, WES = whole-exome sequencing, RNA-seq = RNA sequencing.

## Supplementary Tables

Supplementary Table 1. Patient, treatment, initial, and TIHGG disease characteristics.

Supplementary Table 2. TIHGG treatment and outcomes.

Supplementary Table 3. Methylation-based classification probabilities for the reference dataset for methylation-based classification (Korshunov A and Capper D).

Supplementary Table 4. Methylation-based classification probabilities.

Supplementary Table 5. Methylation-based classification summary.

Supplementary Table 6. Comparison of TIHGG and *de novo* pHGG methylation class.

Supplementary Table 7. Comparison of TIHGG and *de novo* pHGG from the HERBY cohort.

Supplementary Table 8. Chromothripsis evaluation criteria.

Supplementary Table 9. Tier 1 sequence mutations identified from 5 non-hypermutator WES cases without matched germline data.

Supplementary Table 10. Tier 1 sequence mutations identified from 9 non-hypermutator WGS cases with matched germline data.

Supplementary Table 11. TIHGG fusion genes in RNA-seq samples.

Supplementary Table 12. TIHGG structural variations in 10 WGS cases.

Supplementary Table 13. TIHGG copy-number alteration segments in 10 WGS cases.

Supplementary Table 14. Methylation-based classification probabilities for the HERBY dataset

Supplementary Table 15. Germline variants.

Supplementary Table 16. Cytoscape/Metascape results

Supplementary Table 17. GSEA results in TIHGG vs. *de novo* GBM for GO, with all genesets

Supplementary Table 18. GSEA results by category as shown in Fig. 7A.

Supplementary Table 19. Expression Ratio (ER) and p-value (Group B TIHGGs/Group A TIHGGs) for selected genes from GO_DNA_REPAIR geneset.

Supplementary Table 20. *In silico* drug screening results.

Supplementary Table 21. *In vitro* drug screening results.

